# Xanthophyll acyl esters in pepper (*Capsicum annuum*): Delineating the role of two xanthophyll acyltransferases *CaPYP1* and *CaPYP1-like* via gene silencing and genetic complementation

**DOI:** 10.64898/2026.07.28.741266

**Authors:** Jingwei Fu, Bala Rathinasabapathi

## Abstract

Pepper fruit accumulated xanthophyll acyl esters during ripening. To characterize the genes involved, cDNAs for two putative xanthophyll acyltransferases named CaPYP1 and CaPYP1-like were cloned. The deduced amino acid sequences of CaPYP1 and CaPYP1-like had conserved hydrolase and acyltransferase domains. Phylogenetic analyses showed that both putative acyl transferases were conserved in *Viridiplantae*, suggesting important functions for both. CaPYP1 and CaPYP1-like GFP-fusion proteins were localized in the plastid when expressed in leaf tissue. Gene expression analyses indicated that both genes for CaPYP1 and CaPYP1-like were most expressed in ripening fruit (52 to 64 days after anthesis) and senescent leaf, *CaPYP1* having relatively greater expression than *CaPYP1-like*. In virus-induced gene silencing experiments in pepper, xanthophyll esterification was greatly diminished when *CaPYP1* was silenced with a simultaneous change in the ripening fruit’s color from dark red to bright red. In complementation tests, overexpression of *CaPYP1* in a tomato mutant impaired for petal coloration and xanthophyll esterification, resulted in restoration of the petal color and the synthesis of both mono, diacyl and tri esters of xanthophylls. *CaPYP1-like* overexpression in the same genetic background resulted in the synthesis of relatively smaller amounts of xanthophyll monoesters. In an *in vitro* test, zeaxanthin was more sensitive to light than zeaxanthin dipalmitate but both were protected when triacylglycerol was mixed with it, suggesting that acyl moieties could improve xanthophyll stability. Together our results indicate that xanthophyll esterification during pepper fruit ripening is important for fruit color, xanthophyll accumulation and stability and is orchestrated by both CaPYP1 and CaPYP1-like with CaPYP1 playing a major role.

**Highlight:** Ripening pepper fruit accumulates xanthophyll fatty acyl esters associated with nutritional quality and fruit color. The fruit expresses CaPYP1 and CaPYP1-like, two putative acyltransferases in their chromoplasts. In a tomato mutant impaired for xanthophyll esterification, transgenic expression of *CaPYP1* promoted more xanthophyll esterification in ripe fruit than expressing *CaPYP1-like*.

## Introduction

Accumulation of carotenoid pigments in ripe fruit determines the intensity of color, flavors, and nutritional quality of the fruit (Berry et al., 2019). Fruit carotenoids are highly valuable as consumers prefer brightly colored vegetables and fruits (Wang et al., 2025) and provitamin A carotenoids are essential for human health. Carotenoids in flowers and ripening fruits have likely evolved through selective forces via pollinators and seed dispersal agents (Ohmiya et al., 2019). Recent research suggests that apo-carotenoids derived from the breakdown of carotenoids can be signals in modulating plant development and productivity (Rodriguez-Concepcion and Lu, 2026).

Xanthophylls in green tissues have vital roles in protecting the photosynthetic machinery from light damage via the xanthophyll cycle. During fruit ripening, chlorophyll is degraded, chloroplasts are converted to chromoplasts, and xanthophylls and their fatty acyl esters are synthesized and accumulated in plastid microcompartments called plastoglobuli (Ma et al., 2017; Van Wijk, 2017). The esterification of xanthophylls with fatty acids corresponded with increased ripeness and higher levels of accumulation of total carotenoids (Hornero-Mendez and Minguez-Mosquera, 2000). Esterified xanthophylls and free xanthophylls differ in their solubility, thermostability, bioavailability and xanthophyll esters in food need to be hydrolyzed prior to absorption. Hence, research in this area is gaining increased attention from food scientists (Mercadante et al., 2016).

Although ripening fruit of many different taxa including pepper, citrus, mango and peaches are known to accumulate significant levels of xanthophyll esters (Petry and Mercadante, 2016; Ma et al., 2017; Liu et al., 2025), key questions regarding the synthetic pathway to xanthophyll fatty acyl esters are unresolved. These include the identification of genes involved in xanthophyll esterification in ripening fruit and the potential biological advantage for xanthophyll esterification in fruit. In tomato a gene named *PALE YELLOW PETAL 1(PYP1)* when mutated lacked xanthophyll esters in their petals (Ariizumi et al 2014) and the mutant’s chromoplast development was impaired. PYP1 protein contains an α/β hydrolase fold and a lysophosphatidic acyltransferase domain. Lewis et al (2021)’s investigation showed that tomato PYP1 was responsible for xanthophyll esterification in the ripening tomato fruit also. Since tomato fruit expresses PYP1 and its homolog which we named PYP1-like, we named the orthologous proteins expressed in pepper (*Capsicum annuum*) as CaPYP1 and CaPYP1-like respectively. To identify pepper PYP1s involved in xanthophyll acyl ester accumulation in ripening fruit we experimentally tested for the expression of putative *PYP1* genes during fruit ripening, identified their sub-cellular location and whether both CaPYP1 and CaPYP1-like would exhibit the catalytic ability to esterify xanthophylls by using virus-induced gene silencing in pepper and transgenic expression in tomato. Our results elucidate the genetic basis of xanthophyll esterification in pepper fruit and suggest potential biofortification strategies to increase xanthophyll accumulation in fruit vegetables contributing to improved nutritional quality.

## Materials and Methods

### Plant materials

This study used *Capsicum annuum* lines ‘Ruby (line RJ107(6)A3)’, ‘Round of Hungary’ and ‘Early Jalapeño’ from the University of Florida Pepper breeding program, *Solanum lycopersicum* L. lines ‘MicroTom’, and pale yellow petal 1-1 mutant (*pyp1-1(H7L*)) in ‘MicroTom’ background and *Nicotiana benthamiana*. Plants were grown in containers (0.25 to 2 gallon size) using potting medium (Jolly Gardner C25), manually irrigated to container capacity once a day, and supplied with controlled-release fertilizer (Florikan, N:P:K 18:6:8). The growth room had 16-h day length conditions (150 µmol m^-2^ s^-1^) via fluorescent lights and temperature range of 23 to 27°C. When ripe fruit were analyzed, they were harvested after complete ripening as judged based on pericarp color development. For one experiment, fruits at different stages of development were collected based on the number of days after anthesis by tagging individual flowers.

### Sequence comparisons and phylogenetic methods

DNA and amino acid sequences were compared using BLAST (https://blast.ncbi.nlm.nih.gov) and phylogenetic analyses were carried out using Phylogeny (http://www.phylogeny.fr).

### Gene expression

Total RNA was isolated from pepper tissues using RNeasy plant mini kit (Qiagen, Germantown, MD, USA), quantified using a plate reader and the RNA’s integrity was checked by separating on an agarose gel. First-strand cDNA was synthesized using Super-Script III First-strand Synthesis system (Invitrogen, Carlsbad, CA, USA) with random hexamers and oligo(dT) primer (Invitrogen, U.S.A.). The expression levels of *CaPYP1*, *CaPYP1-like*, and *CaCCS* genes (Table S1) were performed by quantitative RT-PCR using green fast qPCR blue mix (AzuzaView^TM^, MA, U.S.A.) with gene-specific primers listed in Table S2. The relative expression levels of targeted genes were quantified using the equation of 2^^–ΔCT^, normalized to the reference gene, encoding ubiquitin-conjugating enzyme E2 (Cheng et al., 2017). Each qPCR reaction was run in three biological replicates and three technical replicates.

### Sub-cellular localization of proteins

Agrobacterium strain GV3101 containing expression constructs were grown in YEP medium containing 50 mg L^−1^ spectinomycin, 10 mg L^−1^ rifampicin, and 20 mg L^−1^ gentamycin at 28°C overnight. In addition, Agrobacterium strain GV3101 containing P19 expression construct (Saxena et al., 2011), was used to promote transient expression of the expressed protein in tobacco leaf. Cells were harvested by centrifugation at 3000 *g* for 15 min at room temperature. The pellet was resuspended in a buffer containing 10 mM MES (pH 5.6), 10 mM MgCl_2_, and 100 mM acetosyringone to a final concentration of OD_600_ of 1.0 each construct. The infiltration solution was prepared by mixing two Agrobacterium strains at a 1:1 ratio, in which one was the pK7FWG2.0 construct and another was the P19 construct. The mixture was infiltrated into expanding leaves of 4-6-week-old *Nicotiana benthamiana* plants using 1-mL syringe without needles. After 2 days, the fluorescent signals of GFP fusion proteins were observed using a confocal microscope (Leica TCS SP5). The argon laser line excitation wavelength and emission bandpass filter wavelengths were 500–550 nm for eGFP fusion proteins and 650–710 nm for chlorophylls. Agrobacterium strain containing GFP only constuct was used as positive control.

### cDNA cloning and Expression Vectors

Protein coding sequences (CDS) of *CaPYP1* and *CaPYP1*-like, minus stop codon were amplified from the cDNA by specific primers listed in Table 2. Protein sequences are listed in Supplementary file 1. The gel-purified PCR products were cloned into an entry vector pENTR™/D-TOPO^®^ (Invitrogen, Waltham, MA, U.S.A.). After validation by Sanger sequencing, XAT cDNAs from the entry vector were subcloned into plant expression vector pK7FWG2.0 by Gateway LR recombination reaction (Invitrogen, U.S.A.) to generate pK7FWG2.0-CaPYP1-eGFP and pK7FWG2.0-CaPYP1-like-eGFP constructs.

### Virus-induced gene silencing

The knock-down pepper lines of *CaPYP1*, *CaPYP1-like*, and *CaCCS* were generated by the VIGS method reported by Kim et al. (2017) with minor modifications as described below. The target region for silencing was designed by the SGN VIGS Tool (https://vigs.solgenomics.net). DNA fragments of 654 bp, 531 bp, and 460 bp of *CaPYP1*, *CaPYP1-like*, and *CaCCS* respectively were amplified from pepper fruit cDNA using specific primers listed with primer names ending in VIGS in Table S2. After subcloning into pTRV2-MCS, which was obtained from Arabidopsis Biological Resource Center (https://abrc.osu.edu/stocks/number/CD3-1040), pTRV2-CaPYP1, pTRV2-CaPYP1-like, and pTRV2-CCS were generated and validated by Sanger sequencing. Each of the vectors was transformed into *Agrobacterium tumefaciens* strain GV3101 by heat shock. The agroinfiltration of pepper plants was performed as previously described by Kim et al. (2017). Briefly, pepper plants with open cotyledons were used for agroinfiltration. *Agrobacterium* strains GV3101 harboring pTRV2 constructs and pTRV1 respectively, were grown overnight at 28°C. After centrifugation and resuspension in infiltration buffer (10 mM MgCl_2_,10 mM MES, 200 mM acetosyringone, pH5.6), densities of both *Agrobacterium* strains were adjusted to 1.0 optical density (OD600 nm) and mixed at ratio 1:1. After incubating at room temperature for 3h, the mixture of *Agrobacterium* strains were infiltrated into abaxial side of cotyledons using 1-mL syringe without needles. The *Agrobacterium* strain GV3101 containing pTRV2-MCS was used as control. Infiltrated plants were kept under dark and high humidity conditions overnight, followed by returning to normal conditions until the reproductive stages.

### Tomato genetic transformation

*Agrobacterium tumefaciens* GV3101 containing each pK7FWG2.0 construct was generated by freeze-thaw transformation and selection on antibiotic-containing agar media plates. Genetic transformation of tomato was achieved using *Agrobacterium tumefaciens* based method modified from (Sun et al., 2006). Briefly, *Agrobacterium* strains harboring constructs were grown in YEP medium containing 50 mg L^−1^ spectinomycin, 10 mg L^−1^ rifampicin, and 20 mg L^−1^ gentamycin at 28 °C overnight. Subsequently, *Agrobacterium* cells were then centrifuged and resuspended in liquid Murashige and Skoog medium (MS medium) (Murashige and Skoog, 1962). *Agrobacterium* cells diluted to 0.5 A600 were used for infection. Surface-sterilized seeds of *pyp1-1* mutant tomato were germinated on half-strength MS (1/2MS) medium. Cotyledons from 10-day old plants were excised and placed on preculture medium [MS medium supplemented with 3% (w/v) sucrose, 2 mg L^−1^ zeatin and 0.8% (w/v) agar] for 2 days. Explants were then immersed in *Agrobacterium* suspensions for 10 min and blotted dry. The infected explants were placed on co-cultivation medium (MS medium supplemented with 3% sucrose, 2 mg L^−1^ zeatin, 100 µM acetosyringone, 10 µM 2-mercaptoethanol and 0.8% agar) at 25°C in the dark for 48h. After co-cultivation, the explants were transferred to the callus induction medium [MS medium supplementing with 3% (w/v) sucrose, 2 mg L^−1^ zeatin, 100 mg L^−1^ kanamycin, 100 mg L^−1^ timentin, 300 mg L^−1^ cefotaxime, and 0.8% (wt./v) agar] for two weeks. Subsequently, the explants were transferred to shoot induction medium (MS medium supplemented with 3% (w/v) sucrose, 1 mg L^−1^ zeatin, 100 mg L^−1^ kanamycin, 100 mg L^−1^ timentin, 300 mg L^−1^ cefotaxime, and 0.8% (wt./v) agar). The shoot induction medium was renewed every two weeks until shoots formed. When the shoots elongated to 2–3 cm, they were excised and transferred to rooting medium [0.5 x MS medium supplemented with 1.5% (w/v) sucrose, 1 mg L^−1^ IAA, 50 mg L^−1^ kanamycin, 100 mg L^−1^ timentin, 300 mg L^−1^ cefotaxime, and 0.6% (w/v) agar]. Rooted plants were transplanted to soil medium. All media were adjusted to pH 5.7 prior to autoclaving. Genomic PCR with T-DNA-specific primers and recording of GFP signal under fluorescence microscopy were performed for identifying the putative transformants. The T_0_ transgenic lines were grown to collect the seeds from ripe fruit, and their T_2_ generations were used for further analysis.

### Carotenoid extraction and Analyses

Carotenoids were extracted from plant tissues using the method of Wahyuni et al. (2011). Precautions were observed to avoid carotenoid loss by exposure to light or oxygen. Saponification of carotenoids was performed based on the method from (Mínguez-Mosquera & Hornero-Méndez, 1993). Briefly, the dried residue of extracted carotenoids was dissolved in 2 mL diethyl ether and 1 ml 10% (w/v) KOH/methanol was added. The material was mixed and reacted for 60 min at room temperature. An aliquot of 3 mL distilled water was added to the mixture, mixed using a vortex mixture and was briefly centrifuged at low speed. The water layer was discarded, and the organic layer was retained and washed with distilled water until it reached neutral pH. An aliquot of 2 mL 2% (w/v) Na_2_SO_4_ solution was used to remove any moisture from the organic layer. The pooled organic solvent was evaporated to dryness under the stream of N_2_. Quantification of carotenoids was performed by HPLC-UV/Vis following the chromatographic method adapted from Delgado-Pelayo & Hornero-Méndez, (2012). The analytic system comprised an Agilent 1100 series chromatograph connected to an Agilent G1314A UV/Vis detector operated by ChemStation software. An Acclaim RSLC 120 C18 column (250 mm x 2.1 mm i.d., 2.2 µm) fitted with a guard column of a similar material (20 mm x 4.6 mm) was used, maintaining the column compartment at 35°C. The N_2_-dried residues of saponified or non-saponified samples were redissolved in acetone prior to HPLC analysis at a flow rate of 0.1 mL/min. The mobile phase consisted of eluent A (water) and eluent B (acetone) using a gradient program as follows: 75% B was increased linearly to 90% B in 20 min, subsequently holding for 15 min, then raised to 95% B in10 min, and maintained for 20 min constantly; finally, eluent B reached 100% in 15 min and maintained for another 20 min. Initial conditions were reached in another 60 min. The carotenoids were monitored at a wavelength of 450 nm, and the spectra were processed using the ChemStation software. Each carotenoid concentration was calculated against the standard curves, which were generated by the same chromatographic method in the concentration ranging from 1–200 ng µL^−1^. The xanthophyll ester fractions were quantified against the calibration curve of zeaxanthin dipalmitate. Concentration was calculated as µg per gram of fresh weight (µg g^−1^ f.wt.).

### Stability of Zeaxanthin and Zeaxanthin dipalmitate under light irradiation

Zeaxanthin and zeaxanthin dipalmitate were purchased from Cayman Chemical (MI, U.S.A.). After dissolving in acetone, 0.5 nmol of zeaxanthin or zeaxanthin dipalmitate mixed with or without mixing with 7 µL of cooking oil (Pure vegetable oil from CRISCO, OH, U.S.A.) were loaded on a silica TLC plate. Subsequently, half part of the plate was exposed to light (100 µmol m^−2^ s^−1^ light intensity supplemented by fluorescent lights) continuously while another part was kept under dark. The plate was photographically recorded at 0, 3, 12, and 24 hours to identify changes in the intensity of colors of the bands.

### LC/(+)ESI mass spectrometry

In an experiment to identify the acyl esters in ripe fruit, pericarp tissue of ‘Jalapeño’ (Jal) and ‘Round of Hungary’ (ROH), were extracted (0.5 g fresh weight in 4.5 ml of methanol:chloroform 2.5:2.0, v/v, mixture containing 0.1% butylated hydroxytoluene). The carotenoid-enriched fraction was dried under a stream of nitrogen and dissolved in 1 mL of ethyl acetate. Ten µL samples of these were analyzed using HPLC UV (450 nm) followed by (+)ESI-mass spectrometry. Instrumentation and analytical conditions were described in Vilarinho et al (2015). Acyl esters of capsanthin, capsorubin, and zeaxanthin were identified based on their HPLC-MS detection, their expected [M+H]^+^ ions with 450-nm peaks in the retention time-window expected of them. Peaks of various carotenoids and acyl carotenoid esters were integrated to evaluate relative levels of various compounds.

### Statistical treatment of quantitative data

The experiments were designed using a completely randomized design with replicate numbers identified in Figure legends. Quantitative data were analyzed using R (version 4.4.x; R Core Team, 2025) by the analysis of variance followed by mean separation tests to identify significant differences between treatment means.

## Results

To understand the relative levels of free carotenoids, free xanthophylls and xanthophyll esters, we analyzed saponified and non-saponified samples of carotenoid extracts from ripe fruit using HPLC/UV, which clearly separated free carotenes, xanthophylls and most monoesters, and diesters (Figure S1). In a preliminary experiment HPLC/(+)ESI-MS was used to identify xanthophyll esters from ripe fruit from two inbred cultivars ‘Round of Hungary’ (ROH), a sweet bell pepper and ‘Early Jalapeno’(JAL), a chili pepper. In this analysis, among free carotenoids, capsanthin was the highest, the next abundant being capsorubin. Monoesters of capsanthin and capsorubin with saturated fatty acyl groups C12, C14 and C16 were identified based on the compounds’ retention time in chromatography and the mass spectral data (Supplementary Figure S2). Peak area analyses were consistent with the hypothesis that the diester forms were less abundant than the monoester forms (Figure S1). In the ripe pepper fruit, zeaxanthin diester forms were less abundant than the corresponding diesters forms of capsanthin and capsorubin (Figure 1).

**Figure. 1.**
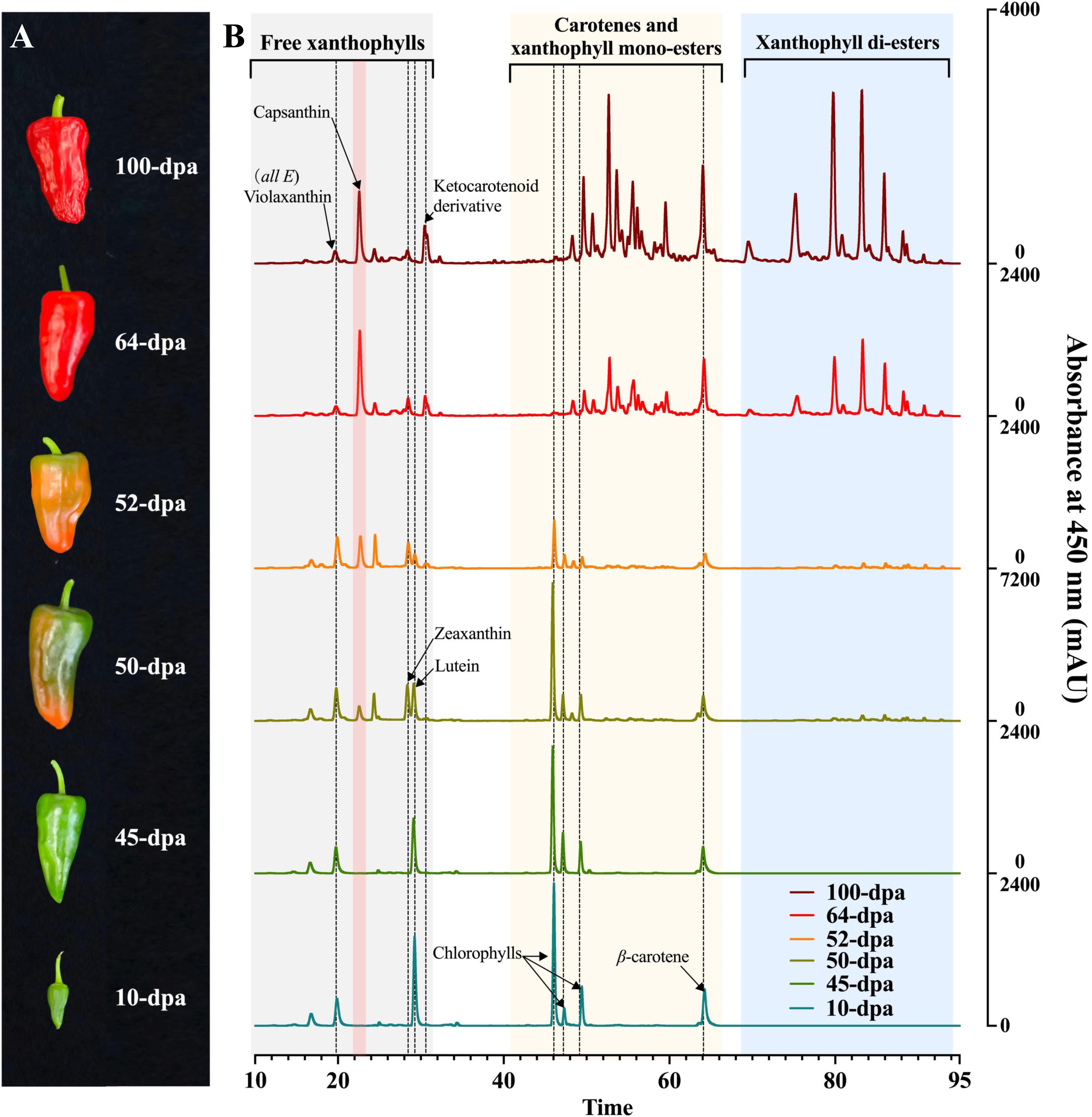
Phenotypes of pepper fruit and their carotenoid profiles. A. The phenotypes of pepper fruit at different developmental and ripening stages. DPA: day post anthesis. B. HPLC chromatograms of carotenoids extracted from pepper fruits at different developmental and ripening stages. HPLC analyses were performed without saponification. The dash-lines across different carotenoid profiles refer to the chromatographic peak of specific carotenoid(s), in corresponding stages of pepper fruit.

Fruits of ‘Ruby’ cultivar were analyzed at 10, 45, 50, 52, 64 and 100 days post anthesis (dpa). Free carotenes, xanthophylls, xanthophyll monoester and diester peaks were identifiable from the time when fruit turned color from green to red (Figure 1), increasing in amounts as the fruit ripened. At 100 dpa, ketocarotenoids capsanthin, capsorubin and a ketocarotenoid derivative were clearly identified (Figure 1). In the ripest fruit more than a dozen xanthophyll ester peaks were identified (Figure 1). When pepper leaves at different developmental stages were examined, xanthophyll ester levels were relatively lower than in fruits and were increased in mature and senescent leaves compared to young leaves (Figure S3)

We cloned two cDNAs orthologous to tomato cDNA for *pyp1* (Ariizumi et al., 2014) from RNA extracted from ripe pepper fruit and named their protein products as CaPYP1 and CaPYP1-like (Table S1). CaPYP1 and CaPYP1-like deduced amino acid sequences were 712 and 675 amino acid residues respectively. Each had a putative plastid-targeting sequence and domains for α/β hydrolase-fold and acyltransferase (Figure S4). Sequence comparisons between these two showed 53% sequence identity and 70% similarity between them. Based on the reference genome, the gene for CaPYP1 was localized in chromosomes 1 of *C. annuum*, *C. chinense*, and *C. baccatum* and the gene for CaPYP1-llike was localized in chromosome 8 of *C. annuum*. Amino acid sequences orthologous to CaPYP1 and CaPYP1-like were identified from multiple plant species to construct a phylogenetic tree shown in Figure S5.

We used a transient expression system in *Nicotiana benthamiana* to test the sub-cellular localization of CaPYP1 and CaPYP1-like proteins. CaPYP1 and CaPYP1-like proteins fused with enhanced green fluorescent protein (GFP) in their C-termini were expressed. GFP fluorescence was co-localized to the plastids for both CaPYP1 and CaPYP1-like (Supplementary Figure S6).

Relative expression levels of *CaPYP1* and *CaPYP1-like* were examined in various vegetative and reproductive tissues of *C. annuum* inbred ‘Ruby’ using quantitative RT-PCR. *CaPYP1* expressed at the highest levels in ripe fruit (52 and 64dpa) with next best expression in senescent leaf with relatively low-level expressions in young fruit, flowers, seed, root, stem and young leaf (Figure 2). Compared to *CaPYP1*, *CaPYP1-like*’s expression was an order of magnitude lower and the most expression was in senescent leaves with next best expression in ripe fruit (Figure 2).

**Figure. 2.**
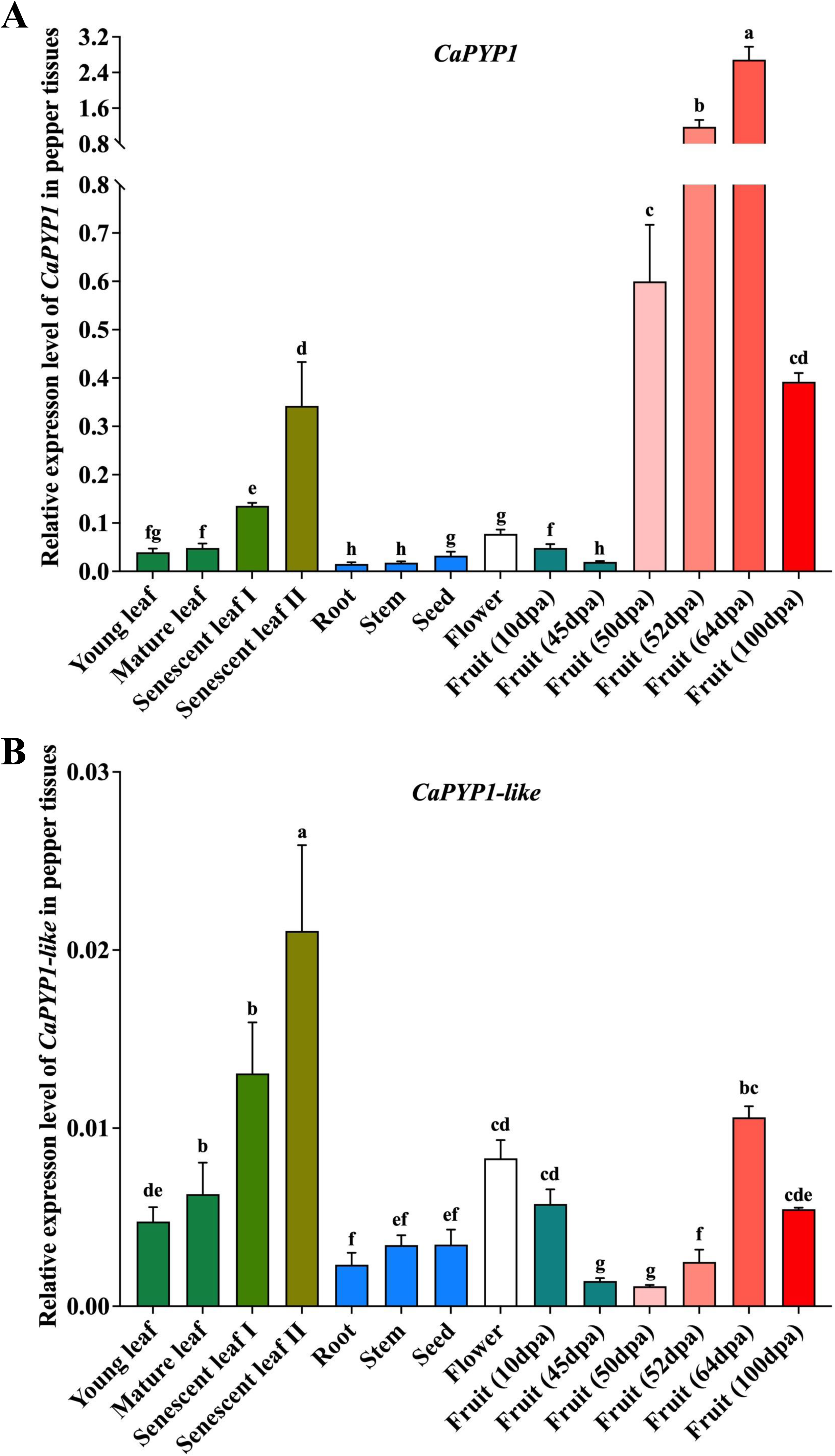
Spatiotemporal expression pattern of *CaPYP1* and *CaPYP1-like* in pepper. The relative expression level of *CaPYP1* (A) and *CaPYP1-like* (B) genes in different pepper tissues. Data of quantitative RT-PCR are presented as mean ± standard deviation of four biological replicates and three technical replicates. Statistical analysis was performed thorough one-way ANOVA and Tukey’s multiple comparisons test. Means significantly different (at *P*=0.05) are marked with different letters.

A virus-induced gene silencing (VIGS) system was used to silence the pepper *CaPYP1* and *CaPYP1-like* singly or in combination. When *CaPYP1* was silenced, the intensity of the red fruit color was reduced than the fruit of the mock control or *CaPYP1-like* silenced fruit (Figure 3aA). When capsanthin/capsorubin synthase (CCS) was silenced, the ripe fruit turned orange (Figure 3A). HPLC-UV analyses of pericarp carotenoid fractions indicated that when *CaPYP1* (singly or in combination with *CaPYP1-like*) was silenced, the levels of xanthophyll monoesters and xanthophyll diesters were significantly reduced including the total carotenoids (Figure 3B and 3C). Quantitative RT-PCR-based relative transcript levels in the pericarp of the silenced fruit indicated that the transcript levels of targeted *PYP1* and *PYP1-like* were significantly reduced compared to the mock and a positive control gene capsanthin/capsorubin synthase (CCS) (Figure 3D to 3F). Similarly, VIGS of CCS reduced the total carotenoids in the pericarp. However, when *CaPYP1-like* was silenced, there was little changes in the carotenoid profiles (Figure 3B) or levels compared to the mock treatment control (Figure 3C).

**Figure 3.**
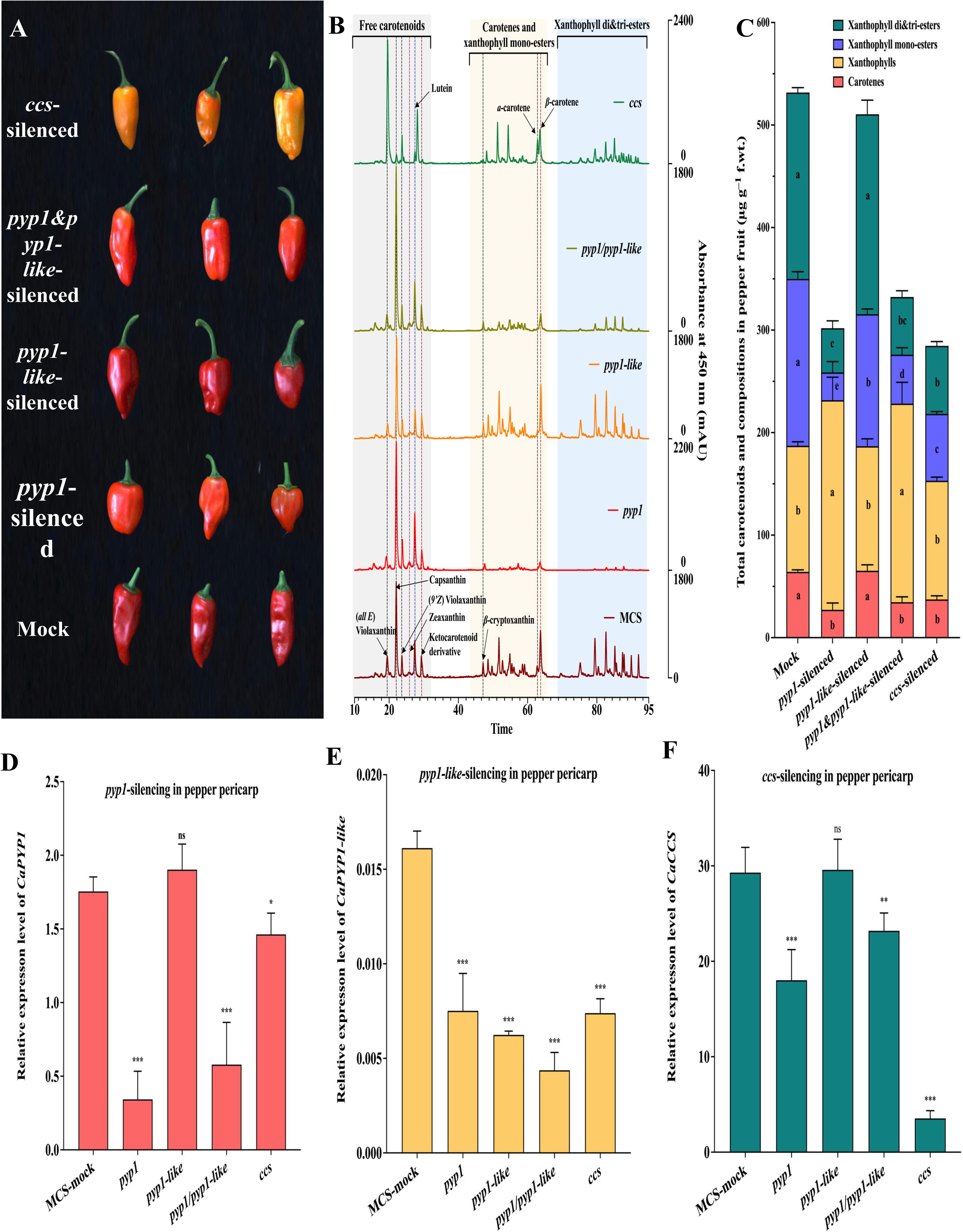
Characterization of *CaPYP1* and *CaPYP1-like* in pepper fruit through the VIGS system. A. The phenotypes of *CaPYP1*-, *CaPYP1-like*-, *CaPYP1 and PYP1-like*, *CaCCS*-silenced and mock pepper fruit. Mock pepper fruit infected with empty TRV-vector and *CaCCS*-silenced pepper fruits were used as negative and positive respectively. Three representative fruits of each treatment were recorded and shown here. B. HPLC chromatograms of *CaPYP1*-, *CaPYP1-like*-, *CaPYP1 & CaPYP1-like*, *CaCCS*-silenced and mock pepper fruits through the VIGS system. HPLC analyses were performed without saponification. C. Total carotenoids and their compositions in pepper fruit (µg g^−1^ fresh weight pericarp). HPLC analyses were performed without saponification. Data were presented as mean ± standard deviation of six biological replicates. Statistical analysis was performed thorough one-way ANOVA and Tukey’s multiple comparisons test. Means significantly different (at *P*=0.05) are marked with different letters. D–F. Relative expression level of *CaPYP1* (D), *CaPYP1-like* (E), and *CaCCS* (F) genes in pepper fruits under VIGS effect. Mean comparisons of target gene-silencing plants against the vector control (mock) were done using the Dunnett test following one-way ANOVA, and results are indicated using asterisks (“*” significant at *P* < 0.05 and “**” significant at *P* < 0.01; ns = not significant). Data of quantitative RT-PCR are presented as mean ± standard deviation of four biological replicates and three technical replicates.

To further test the function of pepper *CaPYP1* and *CaPYP1-like,* we used a genetic complementation approach by generating and characterizing stable transgenic lines of tomato. Tomato mutant defective for *SlPYP1*, *pyp1-1(H7L)*, is characterized by pale yellow petals (Figure 4A). Constitutive expression of *CaPYP1*, restored the petal color to bright yellow in independent transgenic lines (Figure 4A). Transgenic expression of *CaPYP1-like* in *pyp1-1(H7L)* background has also increased the intensity of yellow petal color but less effective than *CaPYP1* expession. Analyses of carotenoids in the flower petals indicated that *pyp1-1(H7L)* had little xanthophyll diesters detectable, transgenic expression of *CaPYP1* increased the levels of free xanthophylls, carotenoids, xanthophyll monoesters and diesters (Figure 4B-F, Figure S7). Transgenic expression of *CaPYP21-like* increased xanthophyll monoesters compared to the control (Figure 4F). Unlike the carotenoid profiles of flowers, analyses of the leaves of the tomato lines indicated no detectable xanthophyll esters even when *CaPYP1* was expressed (Figure S8).

**Figure 4.**
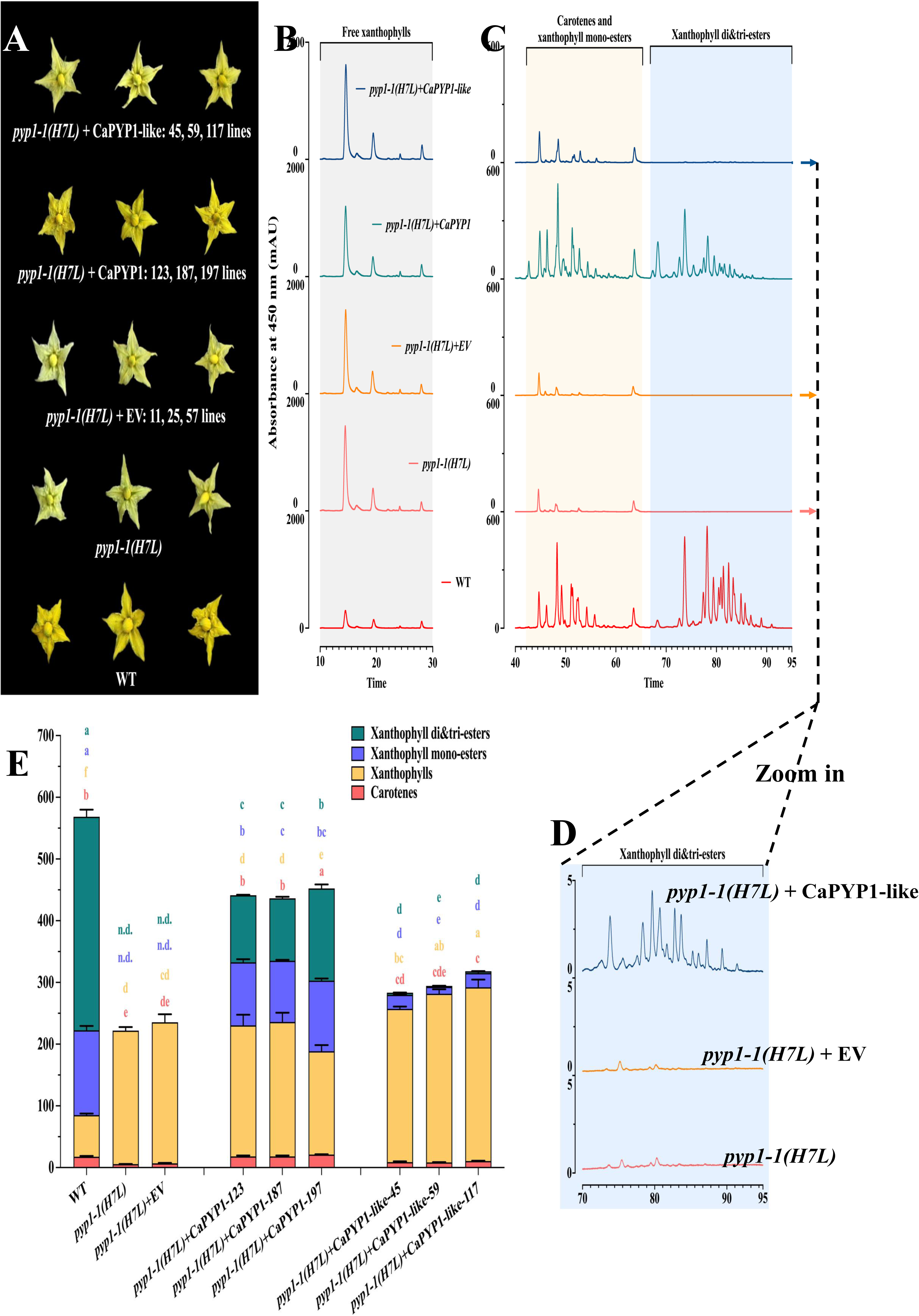
Functional characterization of *CaPYP1* and *CaPYP1-like* through a genetic complementation test. A. Phenotypes of the tomato flowers from independent *CaPYP1* overexpression tomato lines including *CaPYP1*-OE and *CaPYP1-like*-OE in a background of *pyp1-1(H7L)* mutant, empty vector transformed in the same mutant background, *pyp1-1(H7L)* mutant, and WT ‘MicroTom’. Representative flowers from three independent *PYPs*-OE lines are shown here. B&C. The HPLC chromatograms of one representative flower shown in A. D. Zooming-in of HPLC chromatograms from the flower of *pyp1-1(H7L)*, *pyp1-1(H7L)*+EV, and *CaPYP1-like*-OE lines to observe xanthophyll di- and tri-esters if produced. E. Total carotenoids and their compositions in the flowers of three independent *PYP1*-OE and *PYP1-like*-OE lines in *pyp1-1(H7L)* mutant, EV-transformed line, *pyp1-1(H7L)* mutant, and WT ‘MicroTom’. Data of I–L are presented as mean ± standard deviation of a minimum of three biological replicates based on data from HPLC analyses. Statistical analysis was performed thorough one-way ANOVA and Tukey’s multiple comparisons test. Means significantly different (at *P*=0.05) are marked with different letters.

To test the potential function of xanthophyll esterification in improving the stability under light, we compared free xanthophyll to xanthophyll ester under dark or light conditions. When zeaxanthin and zeaxanthin di-palmitate were placed on a silica thin layer plate and exposed to either dark or light conditions for up to 24 h, free zeaxanthin degraded under light by 12 h but zeaxanthin di-palmitate was partially stable up to 12 h (Figure 5). The rate of degradation was reduced when vegetable oil was mixed with xanthophylls (Figure 5).

**Figure 5.**
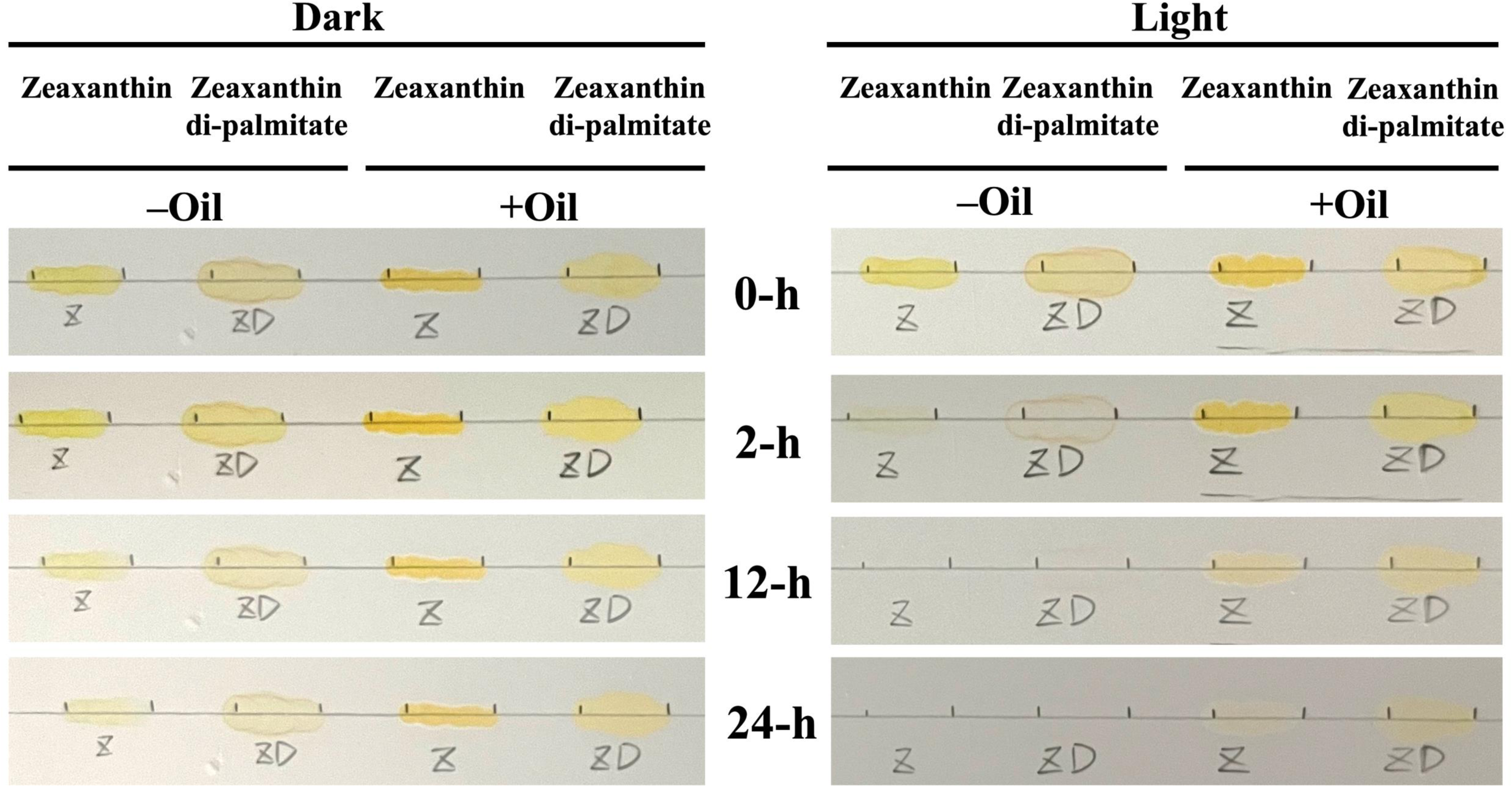
Stability of free xanthophyll and xanthophyll ester under dark and light conditions. Pure zeaxanthin (Z) and zeaxanthin di-palmitate (ZD) were used as free xanthophyll and xanthophyll ester pipetted on to a Silica TLC plate. The same molar concentration of zeaxanthin and zeaxanthin di-palmitate were used in this experiment.

## Discussion

During fruit ripening, degradation of chlorophylls and synthesis and accumulation of carotenoids follow each other (Figure 1, S1 and S2). Xanthophyll acyltransferases expressed during fruit ripening are critical in determining the diversity of xanthophyll fatty acyl esters which are stored in plastoglobuli of the chromoplasts (Li et al., 2023). These enzymes in the esterase/lipase/thioesterase (ELT) protein family are known for xanthophyll esterification in flower petals in transgenic tomato (Fu et al., 2025) but only to a very limited extent in the leaves (Figure S3). But ELT proteins also exhibit multiple other functions including phytyl ester synthesis in senescent leaves (Lippold et al., 2012).

In red ripe peppers, two related ketocarotenoids capsanthin and capsorubin accumulate primarily as their fatty acyl esters (Hornero-Méndez and Mínguez-Mosquera, 2000; Berry et al., 2019). Research studies using animal models have indicated that pepper ketocarotenoids in diet offer multiple health promoting values including cardiovascular protection (Kim et al., 2022), antioxidant and anti-inflammatory activities (Shanmugham and Subban, 2022; Cai et al., 2026) and inhibition of adipogenesis (Jo et al., 2017). Therefore, improving the levels of ketocarotenoid esters in fruits has remained as a key research objective (Ha et al., 2019, Konishi et al., 2019, Fu et al 2026). Despite their importance in nutrition and pepper fruit being an excellent model for xanthophyll ester diversity and accumulation, the genes involved in xanthophyll esterification in ripe peppers have not been characterized.

We report here the identification and experimental validation of two genes involved in xanthophyll esterification in *Capsicum annuum* fruit. Tomato *PYP1*’s role in petal coloration (Ariizumi et al., 2014) and xanthophyll acyltransferase action (Lewis et al., 2021) were used to hypothesize functional roles for Capsicum *CaPYP1* and *CaPYP1-like*. Our hypothesis was supported by the presence of conserved hydrolase fold and acyltransferase domains (Figure S4 and S5) and their expression profiles indicating high level expression in ripening fruit correlating to the accumulation of xanthophyll monoesters and xanthophyll diesters (Figures 1 and 2). Our methodology of splitting carotenoid extracts into two aliquots, saponifying one and not saponifying the other, followed by HPLC-based separation has clearly identified the xanthophyll monoesters and xanthophyll diesters (Figure S1). While the source of fatty acids in the xanthophyll esters in ripening fruit has not been resolved experimentally, we presume this to be degradation of membranes during the conversion of chloroplasts into chromoplasts (Figure S2). Watkin et al (2019) showed that a wheat enzyme catalyzing transesterification of lutein used triacylglycerol as a donor. Additional research is needed on lipid metabolic routes in ripening fruit linked to the synthesis of xanthophyll acyl esters.

Berry et al (2019) showed that fruit color intensity of peppers was correlated with the amounts of capsanthin and capsanthin esters and biosynthesis and accumulation of carotenoids varied in different developmental phases of the fruit. When *CaPYP1* and *CaPYP1-like* expressions were tracked from immature fruit (10 dpa) to ripe stage (64 dpa), both genes’ transcripts increased though *CaPYP1* ‘s expression levels were an order of magnitude greater than that of *CaPYP1-like* (Figure 2). Similarly, expressions of both *CaPYP1* and *CaPYP1-like* genes were progressively increased from young leaves to mature leaves with the highest expression in senescent leaves among the vegetative tissues (Figure 2), suggesting a role for these genes in senescence or low nitrogen status of the leaves (Coulon et al., 2024).

VIGS based gene silencing applied to developing fruit indicated *CaPYP1* plays a major role in the accumulation of total carotenoids, xanthophyll monoesters and xanthophyll diesters (Figure 3). While the fraction of free xanthophylls increased compared to the mock control, the total carotenoid in the silenced fruit significantly decreased (Figure 3) suggesting that xanthophyll esterification improves the flux through the overall carotenoid pathway.

Using the phenotype of ‘pale yellow petal’ allowed us to visually identify the transformants complementing for this phenotype. Flower petal color based selectin has become a convenient system to identify xanthophyll acyltransferases in many different plant taxa (Kishimoto et al., 2020; Zacarías-García et al., 2021; Li et al., 2022, Li et al 2023, Fu et al., 2025). Though *CaPYP1* complemented the mutant (Figure 4), none of the transformants’ flowers expressing the pepper *PYP1* had carotenoid levels equivalent to that of the wild-type suggesting some differences in the substrate preference between tomato native PYP-1 and pepper PYP1.

Mutants for SlPYP1-like is unavailable currently. However, transgenic expression of CaPYP1 in *SlPYP1* mutant background, increased xanthophyll monoesters, diesters and triesters compared to control lines (Figure 4), which is consistent with a hypothesis that CaPYP1 is a functional equivalent of tomato PYP1. Transgenic expression of *CaPYP1-like* also increased the xanthophyll monoesters suggesting the functional overlap between CaPYP1 and CaPYP1-like. Unlike tomato and pepper, *Arabidopsis thaliana* contains a family of ELT proteins whose functional roles are not fully identified (Fu et al. 2025) but suggests that xanthophyll acyl transferases can be potentially useful to engineer ripening fruit to produce specific membrane associated xanthophyll acyl ester compounds with increased stability and biological value.

Although xanthophyll acyl esters are thought to be more stable than the free forms, direct tests for this idea are only a few related to lutein and lutein esters (Mellado-Ortega and Hornero-Méndez. 2017; Metlicar and Albreht, 2022). In this work we noted that zeaxanthin dipalmitate to be more stable under light than zeaxanthin (Figure 5), especially when mixed with oil. Our results establish that *CaPYP1* and *CaPYP1-like* as candidates for xanthophyll esterification during flowering and fruit ripening, a process that stabilizes the flower and fruit coloration. Together we provide foundational tools for improving carotenoid composition, levels, and fruit quality of ripe fruit using breeding and synthetic biology.

In summary, our results identify CaPYP1 as the major xanthophyll acyltransferase responsible for xanthophyll ester formation in ripening pepper fruit, whereas CaPYP1-like exhibits a more limited and partially overlapping function, primarily promoting xanthophyll monoester accumulation. Their plastid localization, ripening-associated expression, and contrasting effects in gene-silencing and tomato complementation assays provide evidence for functional divergence between these two paralogs. The reduction in total carotenoid accumulation following CaPYP1 silencing, together with the greater light stability of zeaxanthin dipalmitate relative to free zeaxanthin, further supports a role for esterification in carotenoid accumulation and pigment stability. These findings clarify the genetic basis of xanthophyll esterification in pepper and identify CaPYP1 as a promising target for breeding and metabolic engineering aimed at improving fruit color, carotenoid retention, and nutritional quality.

## Author contributions

J.F and B.R. designed the study, J.F. planned the experiments, collected data and analyzed them, both authors participated in writing the manuscript and the final version was approved by both authors.

## Data availability

Data are presented in figures and supplementary materials. The primary data that support the findings are available from the corresponding author upon request.

## Acknowledgements

The first author was supported by an assistantship from the Horticultural Sciences Department during his graduate research. We thank Dr. Kari Basso and Dr. Jodie Johnson for their help with mass spectrometry (supported by NIH S10 OD021758-01A1 and S10 ODO30250-01A1) at the Chemistry Department, University of Florida

## Abbreviations

PYP: Pale Yellow Petal
VIGS: Virus induced gene silencing
XAT: Xanthophyll acyltransferase,

## Supplementary Figures

**Figure S1.**
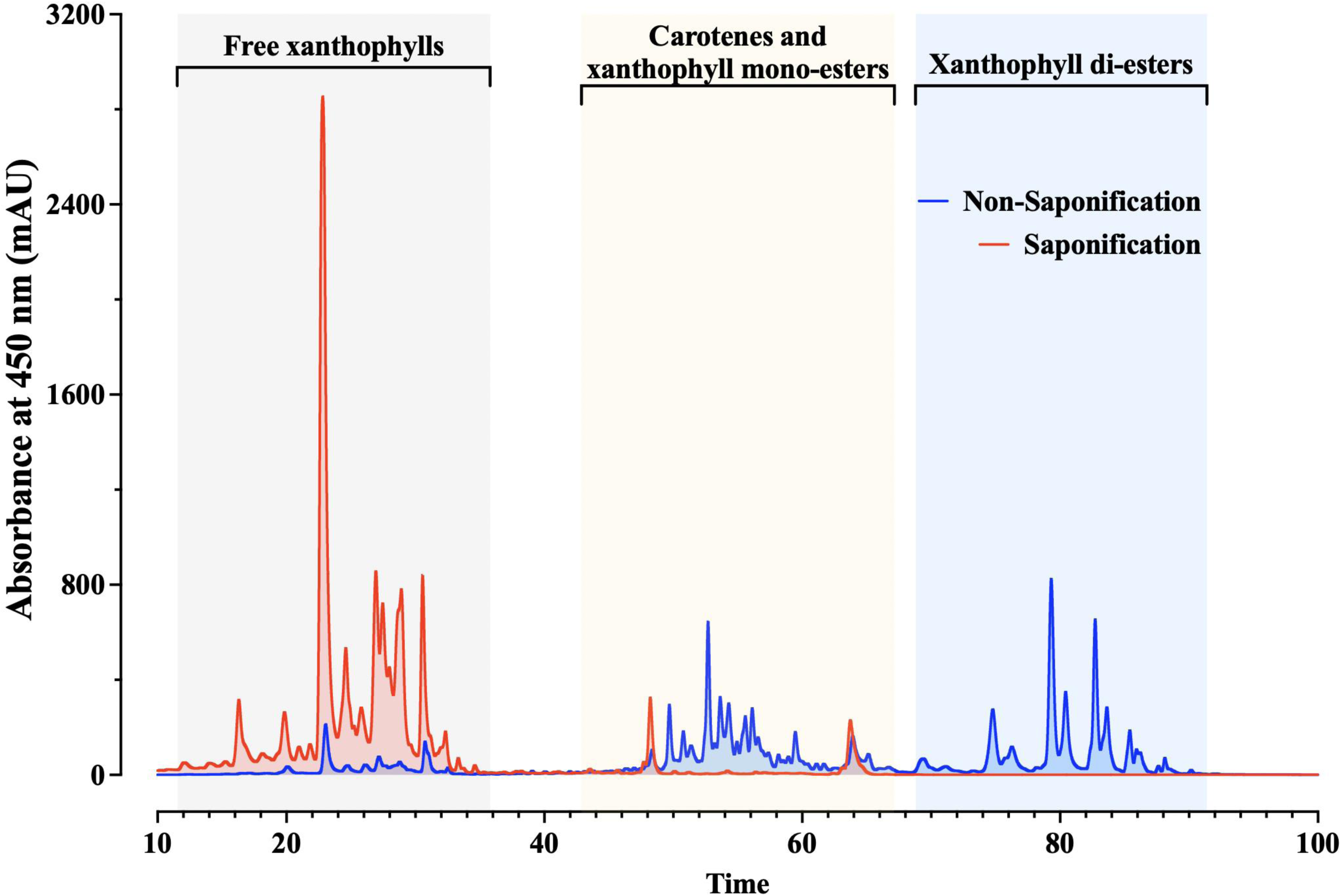
Carotenoid profiles of fully ripe pepper fruit. HPLC chromatograms of carotenoids extracted from fully ripe pepper fruit. HPLC analyses were performed with (in red color) or without saponification (in blue color). The same extract was split into two equal aliquots. One split was saponified with NaOH/MeOH. Another one was kept without saponification. Comparison between unsaponified and saponified extract from pepper fruit showed the conversion from xanthophyll esters to free xanthophylls upon saponification. The xanthophyll esters were detected within regions highlighted with orange and blue background.

**Figure S2.**
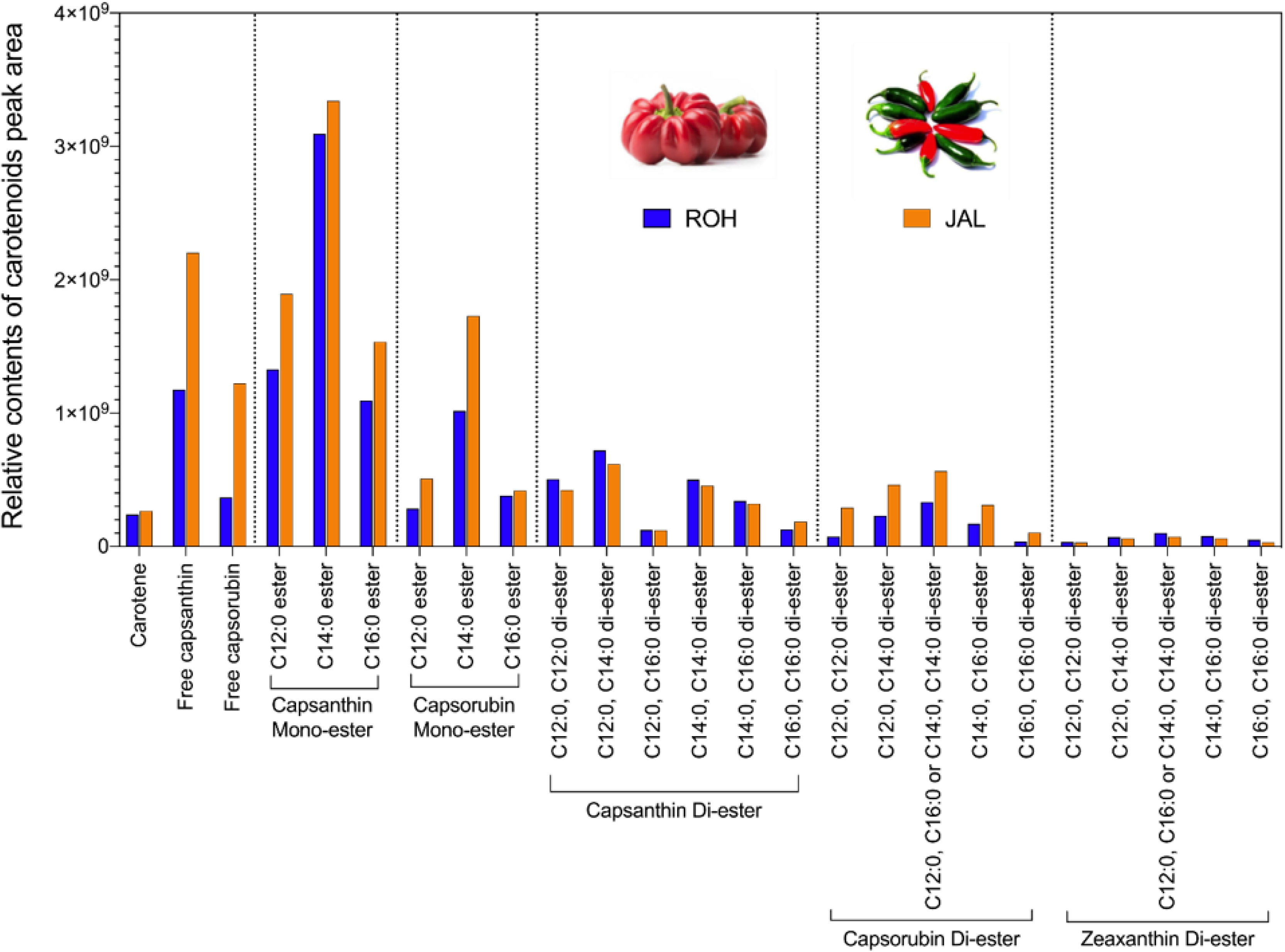
Fractions and relative contents of free carotene, free capsanthin, free capsorubin, and mono and diesters of zeaxanthin, capsanthin and capsorubin in ripe pepper fruit. ROH: ‘Round of Hungary’; JAL: ‘Jalapeno’. The y-axis represents the peak area. To identify the extent of carotenoid esterification in pepper fruit, total carotenoids were extracted from ripe red pericarp of two Capsicum varieties (inset). The extracts were analyzed by HPLC/UV, followed by HPLC/(+)ESI-MS to identify the specific components in each peak. Three replicate extracts were analyzed. Means between ‘ROH’ and ‘JAL’ did not differ significantly using Student’s t-test at *P*=0.05.

**Figure. S3.**
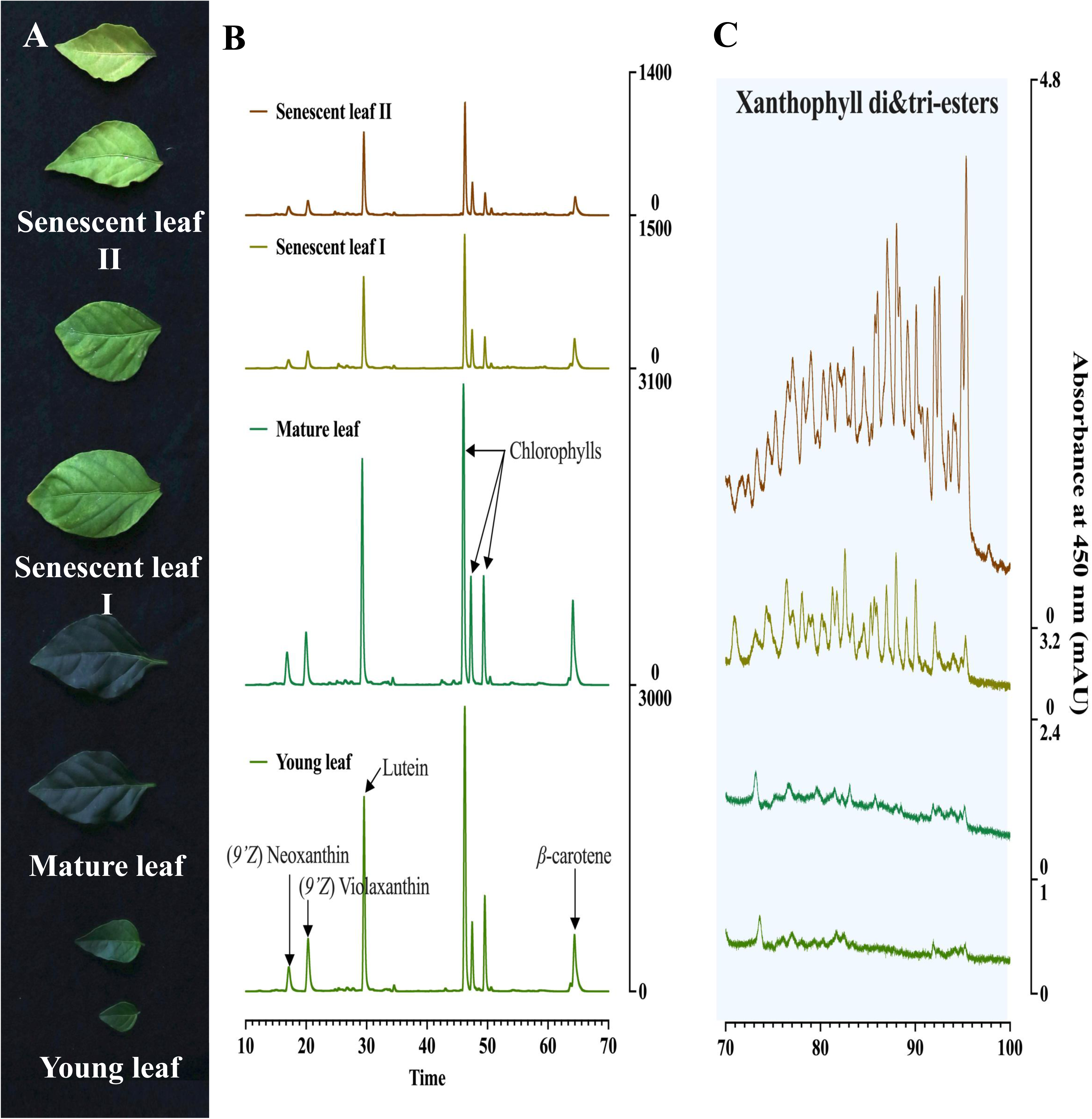
Phenotypes of different types of pepper leaf and their carotenoid profiles. A. The phenotypes of young, fully expanded green, initial senescent, and senescent pepper leaves. Senescent leaf I and II referred to initial and severe senescent pepper leaf, respectively. B&C. Representative HPLC chromatograms of carotenoids extracted from pepper leaf. HPLC analyses were performed without saponification of the carotenoid extracts.

**Figure S4.**
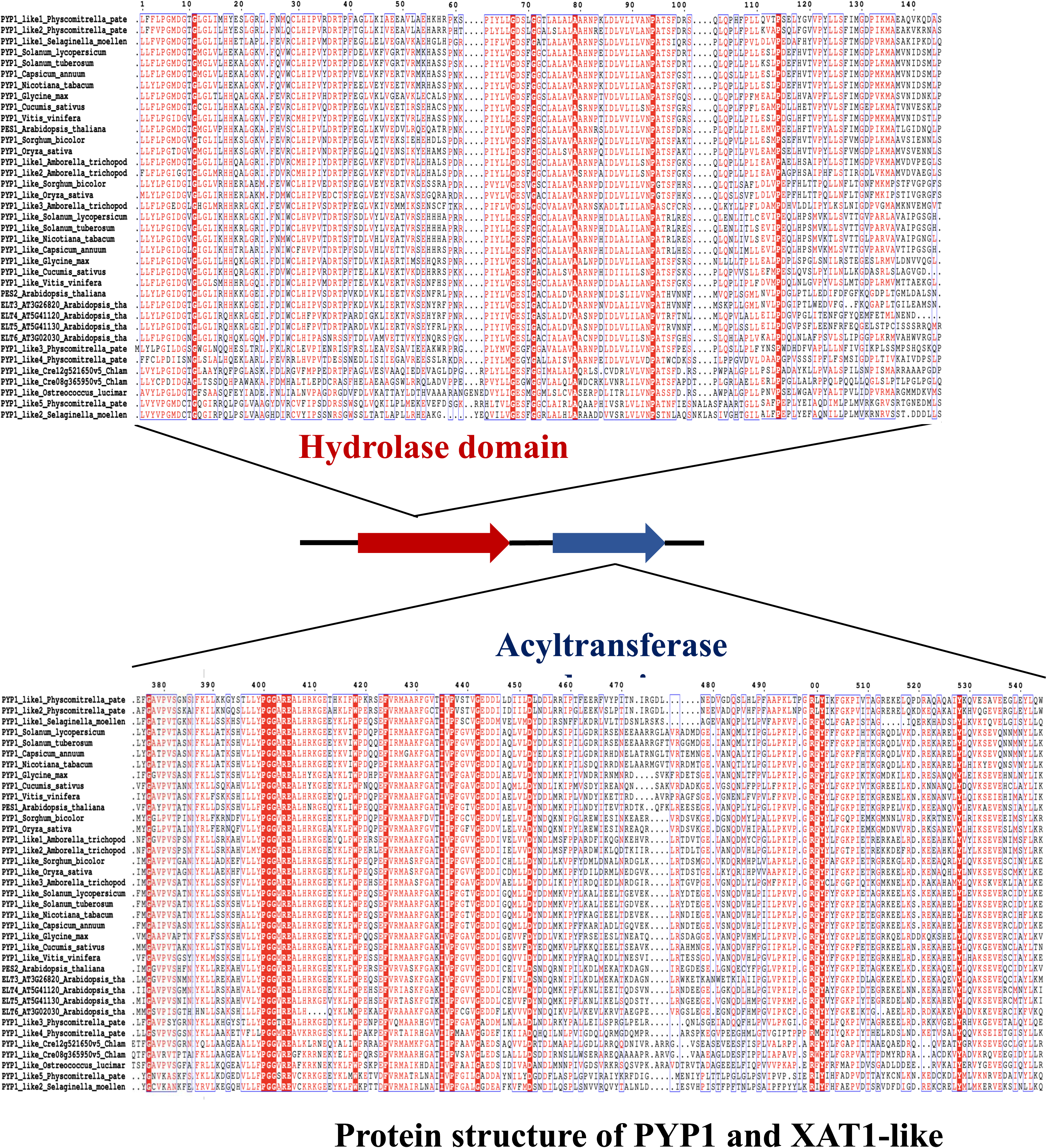
Alignment of PYP1 and its ortholog protein sequences from multiple plant species. Initial alignment of PYP1 and PYP1-like protein sequences was generated using Multiple Sequence Alignment by CLUSTALW (https://www.genome.jp/tools-bin/clustalw). The final alignment was generated using the initial alignment result through ESPript 3.0 (https://endscript.ibcp.fr/ESPript/cgi-bin/ESPript.cgi). Identical amino acid residues from all protein sequences were highlighted in red color. High consensus amino acids from all protein sequences were highlighted in blue box. Low consensus amino acids from all protein sequences were in black color. The accession numbers of the sequences were listed as below: *Capsicum annuum* (CaPYP1, XP_016538484; CaPYP1-like, XP_016561792), *Solanum lycopersicum* (PYP1, XP_004230141; PYP1-like, XP_004231751), *Arabidopsis thaliana* (PES1, AT1G54570; PES2, AT3G2684; ELT3, AT3G26820; ELT4, AT5G41120; ELT5, AT5G41130; ELT6, AT3G02030), *Vitis vinifera* (PYP1, XP_002271452.2; PYP1-like, XP_002274130.1), *Glycine max* (PYP1, XP_003520830.2; PYP1-like, XP_003553970.1), *Cucumis sativus* (PYP1, XP_011648777.1; PYP1-like, XP_004149835.1), *Sorghum bicolor* (PYP1, XP_002455620.1; PYP1-like, XP_021309448.1), *Oryza sativa* (PYP1, XP_015647457.1; PYP1-like, XP_015647440.1), (PYP1, XP_016514310.1; PYP1-like, XP_016488864.1), *Solanum tuberosum* (PYP1, XP_006347827.1; PYP1-like, XP_006338726.1), *Chlamydomonas reinhardtii* (PYP1, Cre12g521650v5), *Physcomitrella patens* (PYP1, XP_024358570.1; PYP1-like, XP_024382778.1; PYP3-like, XP_024388274.1; PYP4-like, XP_024372349.1; PYP5-like, XP_024361534.1). The deduced amino acid sequences contained a hydrolase (in red arrow) and an acyltransferase domain (in blue arrow) by InterPro (Paysan-Lafosse et al., 2023).

**Figure S5.**
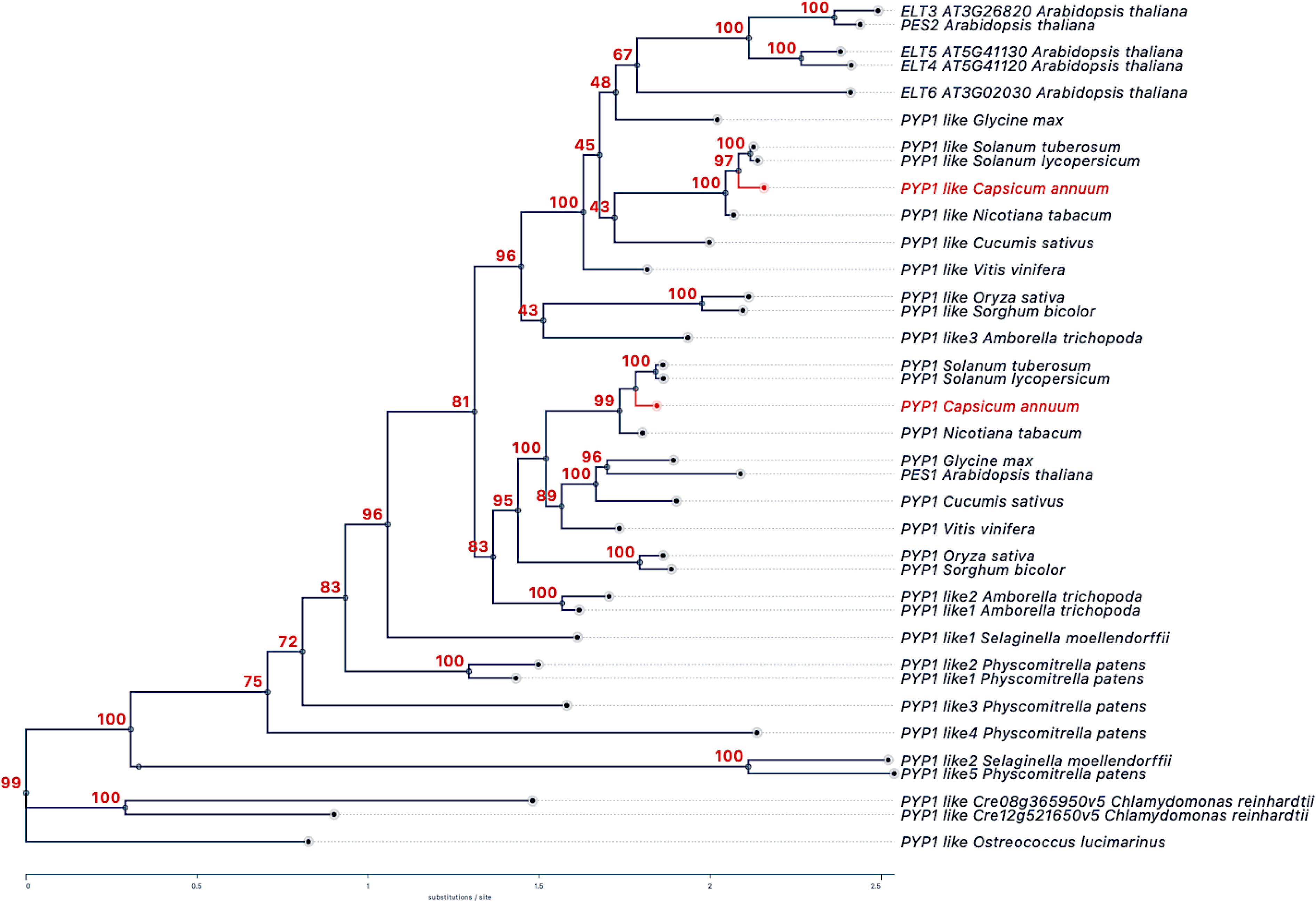
Phylogram of PYP1 and its orthologs from different plant species. Phylogenetic analysis of PYP1 and PYP-like protein sequences from different plant species. The unrooted phylogenetic tree was generated by MEGA 11. The protein sequences were aligned with MUSCLE and then were constructed using the neighbor-joining method. The protein sequences from different plant species were identified through NCBI. The accession number for each protein was the same as protein alignment Fig. S4.

**Figure S6.**
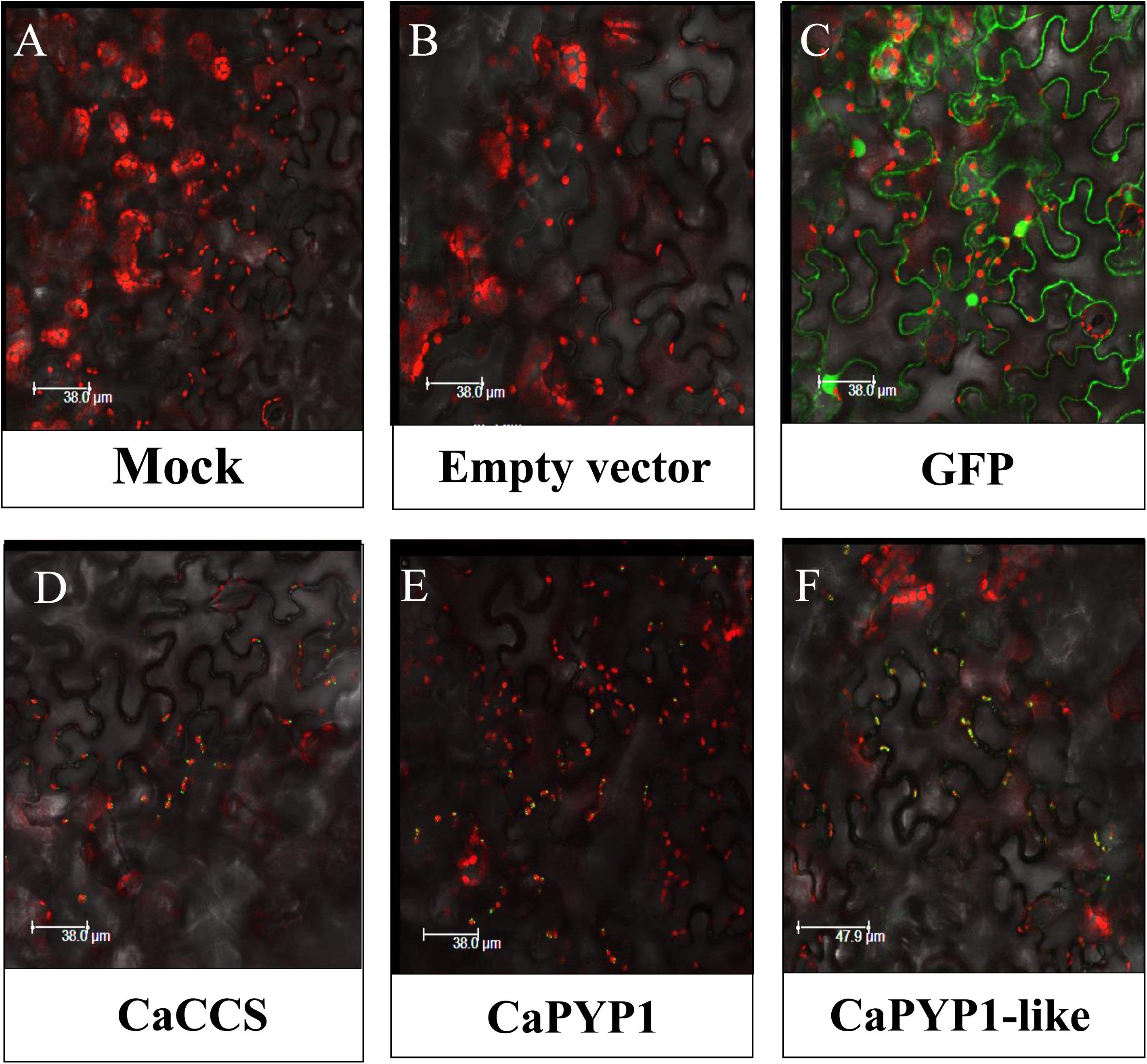
Subcellular localization of PYP1 and PYP1like of tomato and their orthologs from peppers in tobacco leaves. CaPYP1-like, CaPYP2-like, SlPYP1, SlPYP2, and CaCCS were fused to C-terminal eGFP and transiently expressed in tobacco leaves via *Agrobacterium*-mediated transformation. Empty vector and enhanced green fluorescent protein (eGFP) alone was expressed as negative control, which indicated non green signal or a cytosolic and nucleic localization. Representative images were observed under GFP, chlorophyll, and bright field backgrounds and the merged signals are shown.

**Figure S7.**
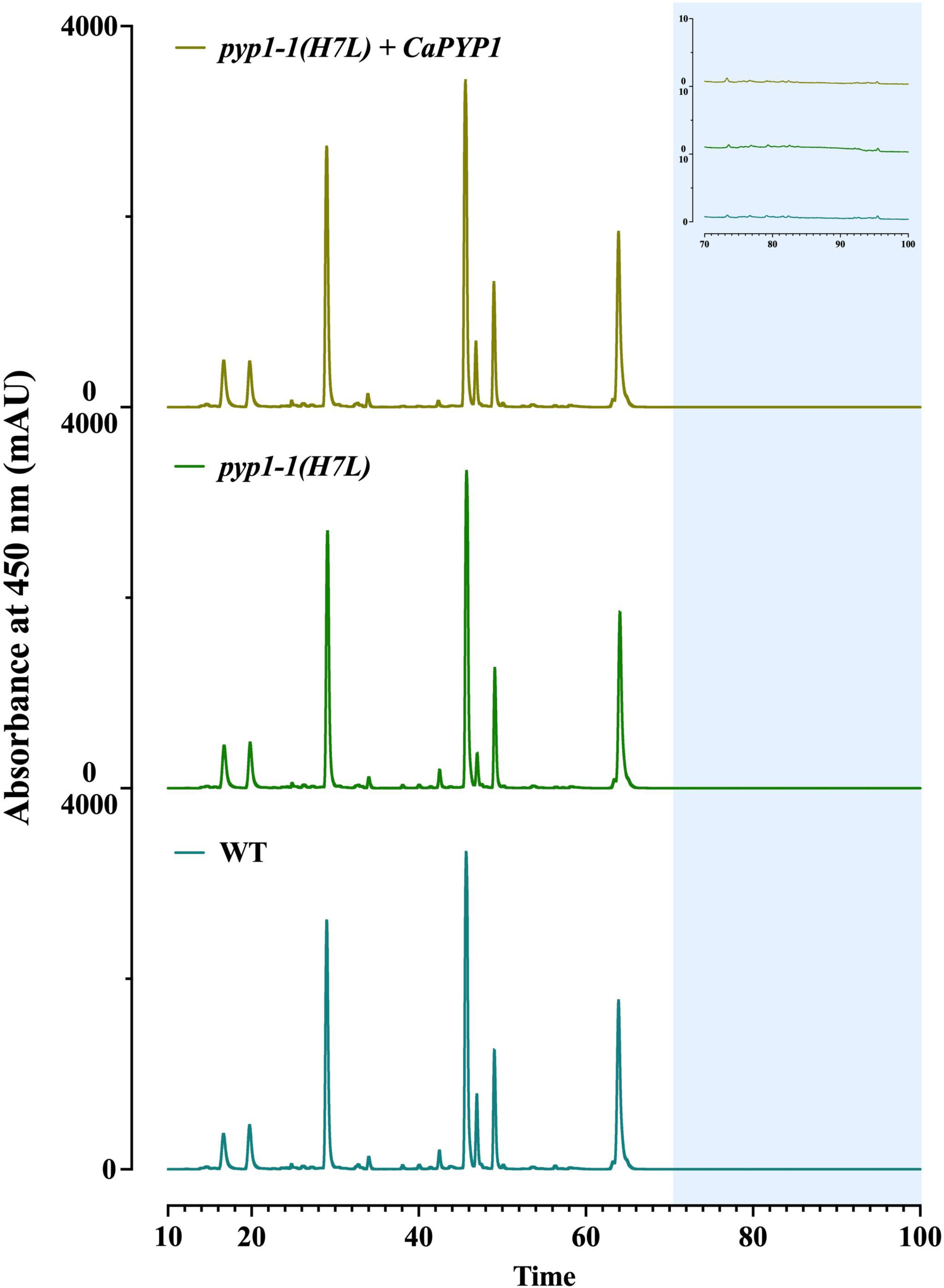
Carotenoid profiles of tomato leaf. HPLC chromatograms of carotenoids extracted from fully expanded mature leaf of WT ‘MicroTom’, *pyp1-1(H7L)*, CaPYP1-OE, and SlPYP1-OE lines,. HPLC analyses were performed without saponification. No xanthophyll esters were detected in green leaf when CaPYP1-like or SlPYP1 was overexpressed in *pyp1-1(H7L)* mutant.

**Figure S8.**
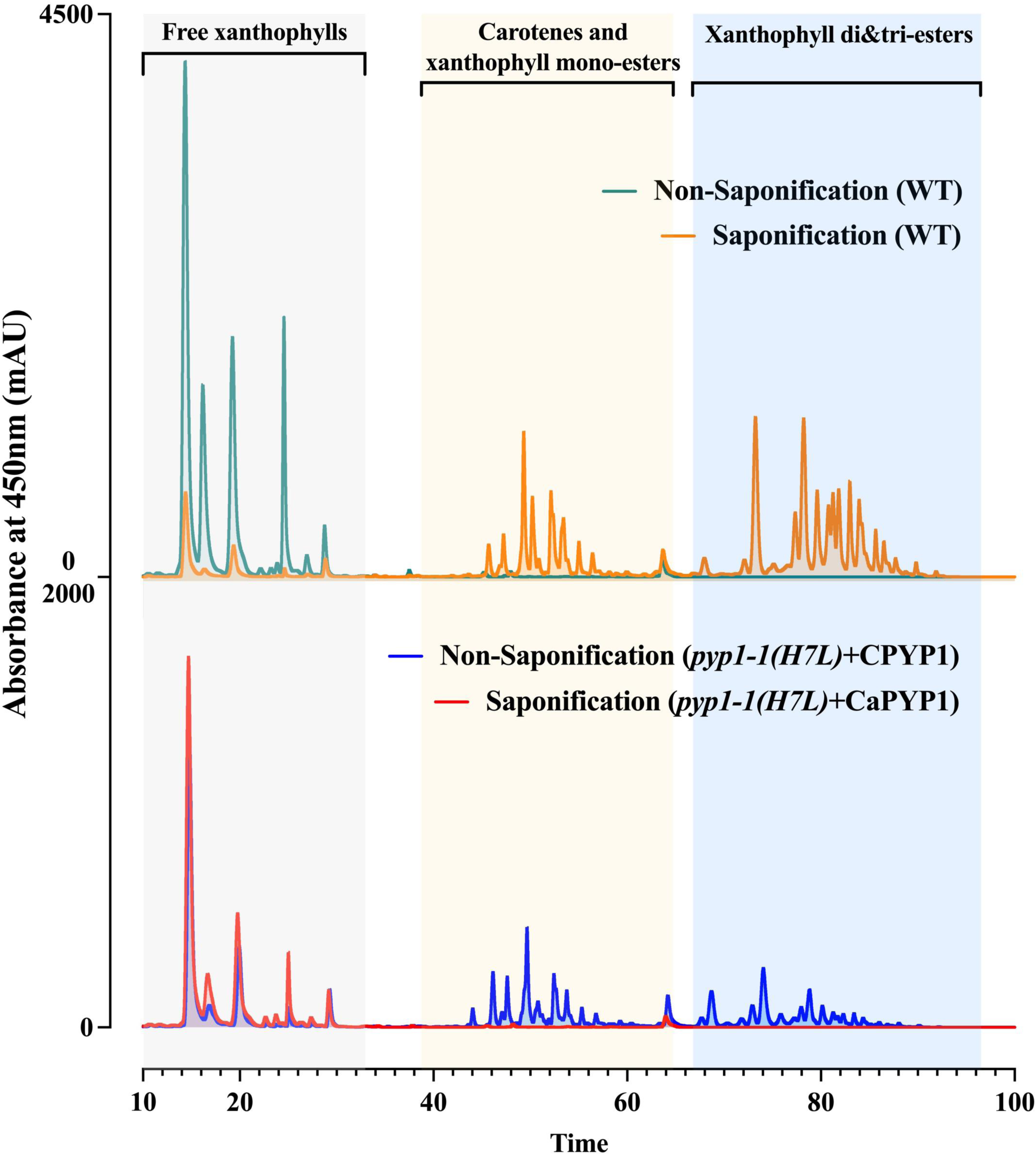
Carotenoid profiles of tomato flowers. HPLC chromatograms of carotenoids extracted from fully open tomato flowers. HPLC analyses were performed with (lines in red or green color line) or without saponification (line in blue or yellow color line). The same extract was split into two equal aliquots. One split was saponified with NaOH/MeOH. Another one had remained without saponification. Comparison between unsaponified and saponified extract from tomato flower showed the conversion from xanthophyll esters to free xanthophylls. The xanthophyll esters were detected within regions highlighted with orange and blue background. The carotenoid profiles with or without saponification of CaPYP1-OE line were similar to the corresponding carotenoid profiles of WT ‘MicroTom’, suggesting the successful biosynthesis of xanthophyll esters in complemented tomato flower.

## Supplementary File 1

### Information on PYP1 and PYP1-like proteins

Tomato PYP-1 XP_004230141(**Chromosome 1, LOC101249584**)

>XP_004230141.1 acyltransferase-like protein At1g54570, chloroplastic isoform X1 [Solanum lycopersicum]

MASLLHNFWAVPRFGLSPDYKPHCIARFACLANRDSTFLSSDSVIVNGVSSIEEKEKSSTIIDVKNSHLA

PAIKEKNKEDIQNKLETLWDDGYGTQTVKDYLEIGSEIIKPDGGPPRWFTPISAGPPLEDSPLLLFLPGM

DGTGMGLVLHEKALGKVFQVWCLHIPVYDRTPFDELVKFVGRTVRMKHASSPNKPIYLVGDSFGGCLALA

IAAHNPEIDLVLILANPATSFDRTQLQPLLPLLESLPDEFHVTVPYLLSFIMGDPLKMAMVNIDSMLPPG

QIIQRLAGNLTDLLAHLYGLADIIPKETLLWKLKLLRSASSYSNSRLHAVNAEVLVIASGKDNMLPSENE

AQRLGNSLRNCTVRYFKDNGHTILLEDGINLLSIIKATSKYRRSKRRDYVKDFLPPSKSEFKNAIKNNSW

YLNITGPVMLSTMENGKIVRGLAGVPREGPVLLVGYHMLMGLEIVPLVQEYMMQTKILLRGIAHPSLFTQ

LVESRPDASSFIDMLKLYGATPVTASNFFKLLATKSHVLLYPGGAREALHRKGEEYKVIWPDQPEFIRMA

AKFGATIVPFGVVGEDDIAQLVLDYDDLKSIPILGDRIRSENEEAARRGLAVRADMDGEIANQMLYIPGL

LPKIPGRFYFFFGKPIHTKGRQDLVKDREKARELYLQVKSEVQNNMNYLLKKREEDPYRNFIDRTMYRAF

SATSGDVPTFDF

Tomato XAT-1-like XP_004231751**(Chromosome 2 LOC101260190)**

>XP_004231751.1 acyltransferase-like protein At3g26840, chloroplastic isoform X1 [Solanum lycopersicum]

MEAISAGTYTAAGVSPLLQSRNSCLRITSARFKIASTAATTGRSTASIGQQPPTVPMSIVDSSELEQTTT

LNMKDFMKWSKDMIASGSEGGPPRWFSPLDCAAPLKDSPLLLYLPGIDGVGLGLIKHHKRLGKIFNIWCL

HVPVTDRTSFSDLVYLVEATVRSEHHHAPRRPIYLLGESFGGCLALAVAARNPHIDLALILANPATRLRE

SQLENLITLCEVIPEQLHPSMVKLLSVTTGVPARLAVAIPGSGHSLQQAVAELFRGDVAFSSYLSVLADV

LPVETLIWRLKILKSAASFVSSRLHAVRSQTLVLSSGKDHLISSPEESEKLHKMLPNCEIRKFNNSGHAL

LLEADFNLVTVIMGANFYRRGRHLDFVRDFVPPSTSEFDSVYQPYRWMEVAFNPVMISTLENGNIVRGLT

GIPSEGPVLLVGYHMMLGLELVPLVSRLWNEHQIVLRGIAHPLMFKKRREGKMPVLSMYDDYRFMGAVPV

SATNFYKLLSSKSHVLLYPGGMREALHRKGEEYKLFWPEQSEFVRMAARFGAKIIPFGTVGEDDIGQMLL

DYDDMMKVPYLKALIEELTGEVEKLRYDTEGEVSNQDVHLPIILPKVPGRFYFYFGKPIETAGRKEELKS

KEKAHELYLEVKSEVERCIDYLKEKRESDSYRNIMARLPYQASHGFDSEVPTFDL

CaPYP-1 (**LOC107839492 Chromosome 8**)

>XP_016538484.1 phytyl ester synthase 1, chloroplastic [Capsicum annuum]

MASFVQTFWAAPRFALTPEYKPRCIARIARLPSRDSTFLPSDSVIVNSVSSIEEREKSSPIADVQNGHLV

STIKEESKKDVQKKLEPLWDDGYGTQTVKDYLEIASEIIKPDGGPPRWFTPISAGPPLEDSPLLLFLPGM

DGTGMGLVLHEKALGKVFQVWCLHIPVYDRTPFVELVKFVERTVRMKHASSPNKPIYLVGDSFGGCLALA

VAAHNPKIDLVLILANPATSFGSTQLQSLLPLLESLPDEFHVTVPYLLSFIMGDPMKMAMVNIDSMLPPG

QISQRLSSNLIDLLVHLSGLADIIPKETLLWKLKLLRSASSYSNSRLHAVNAEVLVLASGKDNMLPSGNE

SQRLKNSLRNCKVRYFKDNGHTILLEDGVNLLSIIKYTNKYRRSKRHDFVMDFVPPSRSEFKNTLKDNRF

YLDCTSPVMLSTMENGKIVRGLAGIPCEGPVLLVGYHMLMGLEIVPLVQEYLRLRKILLRGIAHPTLFTQ

LVESQTNETSFFDTLRLYGATPVSASNFFKLLATKSHVLLYPGGAREALHRKGEEYKVIWPDQQEFIRMG

AKFGATIVPFGVVGEDDIAQLVLDYEDLKSIPILGDRIRHDNELATRNGLTVRTEMNGEVANQALYIPGL

LPKIPGRFYFLFGKPFHTKGRQDLVKDREKARELYLQVKSEVQNNMNYLLKKREEDPYRSVIDRTAYRAF

SATFDDIPTFDY

**Arabidopsis protein corresponding to PYP1: AT1G54570 Gene ‘PES1’**

>NP_564662.1 Esterase/lipase/thioesterase family protein [Arabidopsis thaliana]

MATCSSSLLVLPNLRLSSNQRRNFKVRAQISGENKKATSLEPVNNNGSVSLSTTVQNQKGANEVNGKGKS

KRKIVSDEIELLWDDGYGSKSVKDYFAAAKEILKADGGPPRWFSPVDCGRPVEDAPTLLFLPGMDGTGMG

LVPHHKALGKAFHVSCLHIPVLDRTPFEGLLKVVEDVLRQEQATRPNKPIYLVGDSFGGCLALAVAARNR

SLDLVLILVNPATSFDRSPLQPLLPILEMVPEELHFTVPYALSFIMGDPIKMATLGIDNQLPTGVKIEKL

RQRLTKTMLPLLSELGGIIPRETLLWKLKLLRSGCAYANSRIHAVQAEVLVLASGKDMMLPSQEEAKRLH

GLLKNCSVRCFKDNGHTLLLEDSISLLTVIKGTGKYRRSWRYDLVSDFLPPSKGELAYALDEVLGFLRNA

VGSVFFSTMEDGKIVKGLAGVPDKGPVLLVGYHMLMGLELGPMSEAFIKEKNILFRGMAHPVLYSDNDPA

KAFDYGDWIKVFGAYPVTATNLFKLLDSKSHVLLFPGGAREALHNRGEQYKLIWPEQQEFVRMAARFGAT

IVPFGTVGEDDIAELVLDYNDLMKIPILNDYITEVTRDTKQFKLREESEGEVANQPLYLPGLIPKVPGRF

YYLFGKPIETKGRPELVKDKEEANQVYLEVKAEVENSIAYLLKKREEDPYRSVLDRLNYSLTHTTATHVP

SFEP

XAT2, XP_016561792 CaPYP1-like

Using tomato XAT2 in blast, identified this for putative CaXAT1-like. And it is from CM334 and is on **chromosome 2**:

>PHT92093.1 Acyltransferase-like protein, chloroplastic [Capsicum annuum]

MRMAAISTGVSPCLQYRHFSSTRFSFLVRAASSTASIPHQPPKTPLLSNHRIALEDKTSLIMKGYLEWSK

DMMIADGGPPRWFSPLDCAPPTTIDNFSPLLLYLPGIDGLGIGLIKHHTRLARIFNMWCFHVPVTDRTPF

SELVHLVETTVRSEHHRSPRRPIYLLGESFGGCLALAVAARNPHIDLALILANPATRLHESQLQNLIMLL

EVLPEQLHPSMVKMLSVTTGVPARVAVTIPGSGHPLQQAVAELFRGDVAFSSYLSFEQENDEKETVIYRL

KILKSAAAFVNSRLHAVKAQTLVLSSGKDHLIPSLEESEKLRRMLPNCEIRKFNNSGHVLLLEADFNLVT

VITGANFYRRGRNLDFITDFVPPSTSEFDRVYQPYRWMEVAFNPVMISTLENGNVVRGLTGIPSEGPVLL

VGYHMMLGLELVPLVSSLWYERKIVLRGIAHPLMFKRLREGMMPELSLYDDYRFMGAIPVSASNFYKLLS

SKSHALLYPGGMREALHRKGEEYKLFWPEQSEFVRMAARFGAKIIPFGTVGEDDIGQMLLDYDDLMKIPF

FKARIADLTGQVEKLRNDTEGEVSNQDVHLPIILPKVPGRFYFYFGKPIETEGKKEELKSREKAHELYLE

VKSEVERCLDYLKEKREKDPYRNIMARLPYQATHGFDSQVPTFDL

<u>Protein sequences for CaPYP-1 in other Capsicum species:</u>

*Capsicum frutescens and Capsicum pubescens*

No significant similarities found

*Capsicum chinense* (protein ID 99.72%, 100% query coverage) **(Chromosome 8)**

>PHU09907.1 Acyltransferase-like protein, chloroplastic [Capsicum chinense]

MASFVQTFWAAPRFALTPEYKPRCIARIGRLPSRDSTFLPSDSVIVNSVSSIEEREKSSPIADVQNGHLV

STIKEESKKDVQKKLEPLWDDGYGTQTVKDYLEIASEIIKPDGGPPRWFTPISAGPPLEDSPLLLFLPGM

DGTGMGLVLHEKALGKVFQVWCLHIPVYDRTPFVELVKFVERTVRMKHASSPNKPIYLVGDSFGGCLALA

VAAHNPKIDLVLILANPATSFGSTQLQSLLPLLESLPDEFHVTVPYLLSFIMGDPMKMAMVNIDSMLPPG

QISQRLSSNLIDLLVHLSGLADIIPKETLLWKLKLLRSASSYSNSRLHAVNAEVLVLASGKDNMLPSGNE

SQRLKNSLRNCKVRYFKDNGHTILLEDGVNLLSIIKYTNKYRRSKRHDFVMDFVPPSRSEFKNTLKDNRF

YLDCTSPVMLSTMENGKIVRGLAGIPCEGPVLLVGYHMLMGLEIVPLVQEYLRQRKILLRGIAHPTLFTQ

LVESQTNETSFFDTLRLYGATPVSASNFFKLLATKSHVLLYPGGAREALHRKGEEYKVIWPDQQEFIRMG

AKFGATIVPFGVVGEDDIAQLVLDYEDLKSIPILGDRIRHDNELATRNGLTVRTEMNGEVANQALYIPGL

LPKIPGRFYFLFGKPFHTKGRQDLVKDREKARELYLQVKSEVQNNMNYLLKKREEDPYRSVIDRTAYRAF

SATFDDIPTFDY

*Capsicum baccatum* (**Chromosome 8 MLFT02000008.1**)

>PHT41226.1 Acyltransferase-like protein, chloroplastic [*Capsicum baccatum*]

MASFVQTFWAAPRFALTPEYKPRCIARIARLPSRDSTFLPSDSVIVNSVSSIEEREKSSPIADVQNGHLV

STIKEESKKDVQKKLEPLWDDGYGTQTVKDYLEIASEIIKPDGGPPRWFTPISAGPPLEDSPLLLFLPGM

DGTGMGLVLHEKALGKVFQVWCLHIPVYDRTPFVELVKFVERTVRMKHASSPNKPIYLVGDSFGGCLALA

VAAHNPKIDLVLILANPATSFGSTQLQSLLPLLESLPDEFHVTVPYLLSFIMGDPMKMAMVNIDSMLPPG

QIIQRLSSNLIDLLVHLSGLADIIPKETLLWKLKLLRSASSYSNSRLHAVNAEVLVLASGKDNMLPSGNE

SQRLKNSLRNCKVRYFKDNGHTILLEDGVNLLSIIKYTNKYRHSKRHDFVMDFVPPSRSEFKNALKDNRF

YLDCTSPVMLSTMENGKIVRGLAGIPCEGPVLLVGYHMLMGLEIVPLVQEYLRLRKILLRGIAHPTLFTQ

LVESQTNETSFFDTLRLYGATPVSASNFFKLLATKSHVLLYPGGAREALHRKGEEYKVIWPDQQEFIRMG

AKFGATIVPFGVVGEDDIAQLVLDYEDLKSIPILGDRIRHDNELATRNGLTVRTEMNGEVANQALYIPGL

LPKIPGRFYFLFGKPIHTKGRQDLVKDREKARELYLQVKSEVQNNMNYLLKKREEDPYRSVIDRTAYRAF

SATFDDIPTFDY

When CaPYP1-like sequence was used as a query, only PYP-1’s turned up with significant homology in *C. chinense* and *C. baccatum.*

*Capsicum frutescens and Capsicum pubescens*

No significant similarities found.

*Capsicum chinense*

>PHU09907.1 Acyltransferase-like protein, chloroplastic [Capsicum chinense]

MASFVQTFWAAPRFALTPEYKPRCIARIGRLPSRDSTFLPSDSVIVNSVSSIEEREKSSPIADVQNGHLV

STIKEESKKDVQKKLEPLWDDGYGTQTVKDYLEIASEIIKPDGGPPRWFTPISAGPPLEDSPLLLFLPGM

DGTGMGLVLHEKALGKVFQVWCLHIPVYDRTPFVELVKFVERTVRMKHASSPNKPIYLVGDSFGGCLALA

VAAHNPKIDLVLILANPATSFGSTQLQSLLPLLESLPDEFHVTVPYLLSFIMGDPMKMAMVNIDSMLPPG

QISQRLSSNLIDLLVHLSGLADIIPKETLLWKLKLLRSASSYSNSRLHAVNAEVLVLASGKDNMLPSGNE

SQRLKNSLRNCKVRYFKDNGHTILLEDGVNLLSIIKYTNKYRRSKRHDFVMDFVPPSRSEFKNTLKDNRF

YLDCTSPVMLSTMENGKIVRGLAGIPCEGPVLLVGYHMLMGLEIVPLVQEYLRQRKILLRGIAHPTLFTQ

LVESQTNETSFFDTLRLYGATPVSASNFFKLLATKSHVLLYPGGAREALHRKGEEYKVIWPDQQEFIRMG

AKFGATIVPFGVVGEDDIAQLVLDYEDLKSIPILGDRIRHDNELATRNGLTVRTEMNGEVANQALYIPGL

LPKIPGRFYFLFGKPFHTKGRQDLVKDREKARELYLQVKSEVQNNMNYLLKKREEDPYRSVIDRTAYRAF

SATFDDIPTFDY

>PHT41226.1 Acyltransferase-like protein, chloroplastic [*Capsicum baccatum*]

MASFVQTFWAAPRFALTPEYKPRCIARIARLPSRDSTFLPSDSVIVNSVSSIEEREKSSPIADVQNGHLV

STIKEESKKDVQKKLEPLWDDGYGTQTVKDYLEIASEIIKPDGGPPRWFTPISAGPPLEDSPLLLFLPGM

DGTGMGLVLHEKALGKVFQVWCLHIPVYDRTPFVELVKFVERTVRMKHASSPNKPIYLVGDSFGGCLALA

VAAHNPKIDLVLILANPATSFGSTQLQSLLPLLESLPDEFHVTVPYLLSFIMGDPMKMAMVNIDSMLPPG

QIIQRLSSNLIDLLVHLSGLADIIPKETLLWKLKLLRSASSYSNSRLHAVNAEVLVLASGKDNMLPSGNE

SQRLKNSLRNCKVRYFKDNGHTILLEDGVNLLSIIKYTNKYRHSKRHDFVMDFVPPSRSEFKNALKDNRF

YLDCTSPVMLSTMENGKIVRGLAGIPCEGPVLLVGYHMLMGLEIVPLVQEYLRLRKILLRGIAHPTLFTQ

LVESQTNETSFFDTLRLYGATPVSASNFFKLLATKSHVLLYPGGAREALHRKGEEYKVIWPDQQEFIRMG

AKFGATIVPFGVVGEDDIAQLVLDYEDLKSIPILGDRIRHDNELATRNGLTVRTEMNGEVANQALYIPGL

LPKIPGRFYFLFGKPIHTKGRQDLVKDREKARELYLQVKSEVQNNMNYLLKKREEDPYRSVIDRTAYRAF

SATFDDIPTFDY

## Tables

**Table S1.** Accession numbers of PYP1, PYP1-like and CaCCS gene sequences.

| Gene | Plant species | Accession number | Accession (NCBI) |
| --- | --- | --- | --- |
| CaPYP1 | *Capsicum annuum* L. cv. Ruby | Capana08g001343 or CA08g11060 | XP_016538484 |
| CaPYP1-like | *Capsicum annuum* L. cv. Ruby | Capana02g003704 or CA02g31200 | XP_016561792 |
| CaCCS | *Capsicum annuum* L. cv. Ruby | Capana06g000615 or CA06g22860 | NP_001311998 |

**Table S2.**
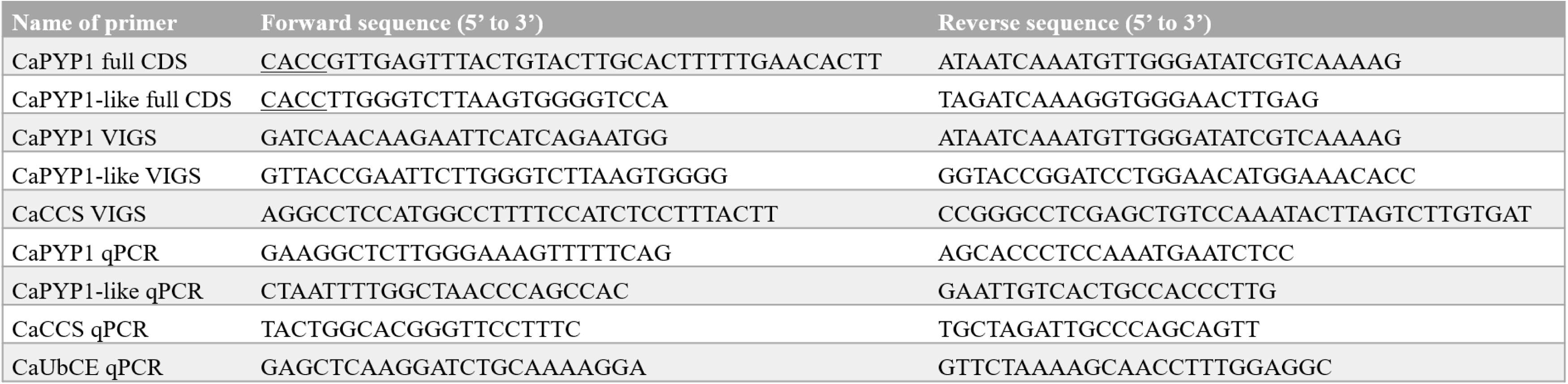
List of primers used in this study. Primers named with ‘full CDS’ were used for amplifying the full length cDNAs, those with ‘VIGS’ were used for cloning DNA used for the silencing experiment, those with ‘qPCR’ were used for gene expression studies.

